# Decoding the genomic basis of osteoarthritis

**DOI:** 10.1101/835850

**Authors:** Julia Steinberg, Lorraine Southam, Natalie C Butterfield, Theodoros I Roumeliotis, Andreas Fontalis, Matthew J Clark, Raveen L Jayasuriya, Diane Swift, Karan M Shah, Katherine F Curry, Roger A Brooks, Andrew W McCaskie, Christopher J. Lelliott, Jyoti S Choudhary, JH Duncan Bassett, Graham R Williams, J Mark Wilkinson, Eleftheria Zeggini

**Affiliations:** Institute of Translational Genomics, Helmholtz Zentrum München – German Research Center for Environmental Health, 85764 Neuherberg, Germany; Cancer Research Division, Cancer Council NSW, Sydney, New South Wales 2011, Australia; Wellcome Sanger Institute, Hinxton CB10 1SA, United Kingdom; Molecular Endocrinology Laboratory, Department of Metabolism, Digestion and Reproduction, Imperial College London, London W12 ONN, UK; The Institute of Cancer Research, London SW7 3RP, UK; Department of Oncology and Metabolism, University of Sheffield, Sheffield S10 2RX, UK; Division of Trauma & Orthopaedic Surgery, Department of Surgery, University of Cambridge, Cambridge CB2 2QQ, UK; Centre for Integrated Research into Musculoskeletal Ageing and Sheffield Healthy Lifespan Institute, University of Sheffield, Sheffield S10 2TN, UK

**Keywords:** Osteoarthritis, translational genomics, drug targets, drug repurposing, electronic health record, RNA sequencing, proteomics, patient stratification, functional genomics

## Abstract

Osteoarthritis causes pain and functional disability for a quarter of a billion people worldwide, with no disease-stratifying tools nor modifying therapy. Here, we use primary cartilage and synovium from osteoarthritis patients to construct a molecular quantitative trait locus map of gene expression and protein abundance. By integrating data across omics levels, we identify likely effector genes for osteoarthritis-associated genetic signals. We detect pronounced molecular differences between macroscopically intact and highly degenerated cartilage. We identify molecularly-defined patient subgroups that correlate with clinical characteristics, stratifying patients on the basis of their molecular profile. We construct and validate a 7-gene classifier that reproducibly distinguishes between these disease subtypes, and identify potentially actionable compounds for disease modification and drug repurposing.

---

Osteoarthritis is a severe, debilitating disease, affecting ̃240 Million people worldwide^1^. It is a heterogeneous disease^2^, hallmarked by cartilage degeneration and synovial hypertrophy. The lifetime risk of developing symptomatic knee and hip osteoarthritis is estimated to be 45% and 25%, respectively^3, 4^, and is on an upward trajectory commensurate with rises in obesity and the ageing population. Older age, female sex, obesity, joint morphology and injury are established clinical risk factors for osteoarthritis, and genome-wide association studies (GWAS) have identified ̃90 robustly-replicating risk loci^5^.

There is no cure for osteoarthritis. Disease management focusses on alleviating pain, and in end-stage disease the only treatment is joint replacement surgery. Two million arthroplasties are carried out annually in the European Union alone (http://appsso.eurostat.ec.europa.eu/nui/show.do?dataset=hlth_co_proc2&lang=en), emphasising the clear and urgent need to develop new therapies that alter the natural history of disease rather than deal with its consequences. To achieve this, we need to improve our understanding of the underlying molecular mechanisms of osteoarthritis pathogenesis and of progression from low-to high-grade disease. Successful future treatment needs to reflect the heterogeneity of osteoarthritis and requires the identification of biological endotypes to which relevant therapeutic modalities may be tailored.

In osteoarthritis, we are able to access the disease-relevant tissue at the point of surgery, allowing the study of paired *ex vivo* samples from as yet unaffected and of highly degenerated tissue at the affected organ. Through in-depth genomics characterisation of human primary tissue collected from patients undergoing joint replacement, we are able to significantly enhance our understanding of disease processes, identify likely effector genes for hitherto unresolved genetic association signals, and move towards precision medicine by building classifiers for patient stratification based on molecular signatures.

## Results

### Molecular map of primary tissue

Here, to improve our understanding of the molecular profile of key osteoarthritis cell types, we collected low-grade (macroscopically intact) and high-grade (highly degraded) cartilage, and synovial tissue samples from 115 patients undergoing joint replacement for osteoarthritis. All cartilage samples were collected from weight-bearing areas of the joint to ensure that any differences observed between low- and high-grade cartilage reflect disease progression stage rather than differential biomechanical stress. All three tissues were profiled by RNA sequencing, and cartilage samples were also profiled by quantitative proteomics (Figure 1). After quality control, we assessed the expression of 15,249 genes in cartilage and 16,004 genes in synovium. We detected and quantified the abundance of 1,677 proteins across all patients, and of 4,801 proteins in at least 30 patients, in line with the resolution depth of the isobaric labelling method employed. We generated genome-wide genotype data from peripheral blood, imputing to 10,249,108 autosomal sequence variants to identify molecular quantitative trait loci (QTLs) in each tissue and omics type. This provides a first in-depth map of genetically-determined gene and protein level regulation in osteoarthritis-relevant tissues.

**Figure 1.**
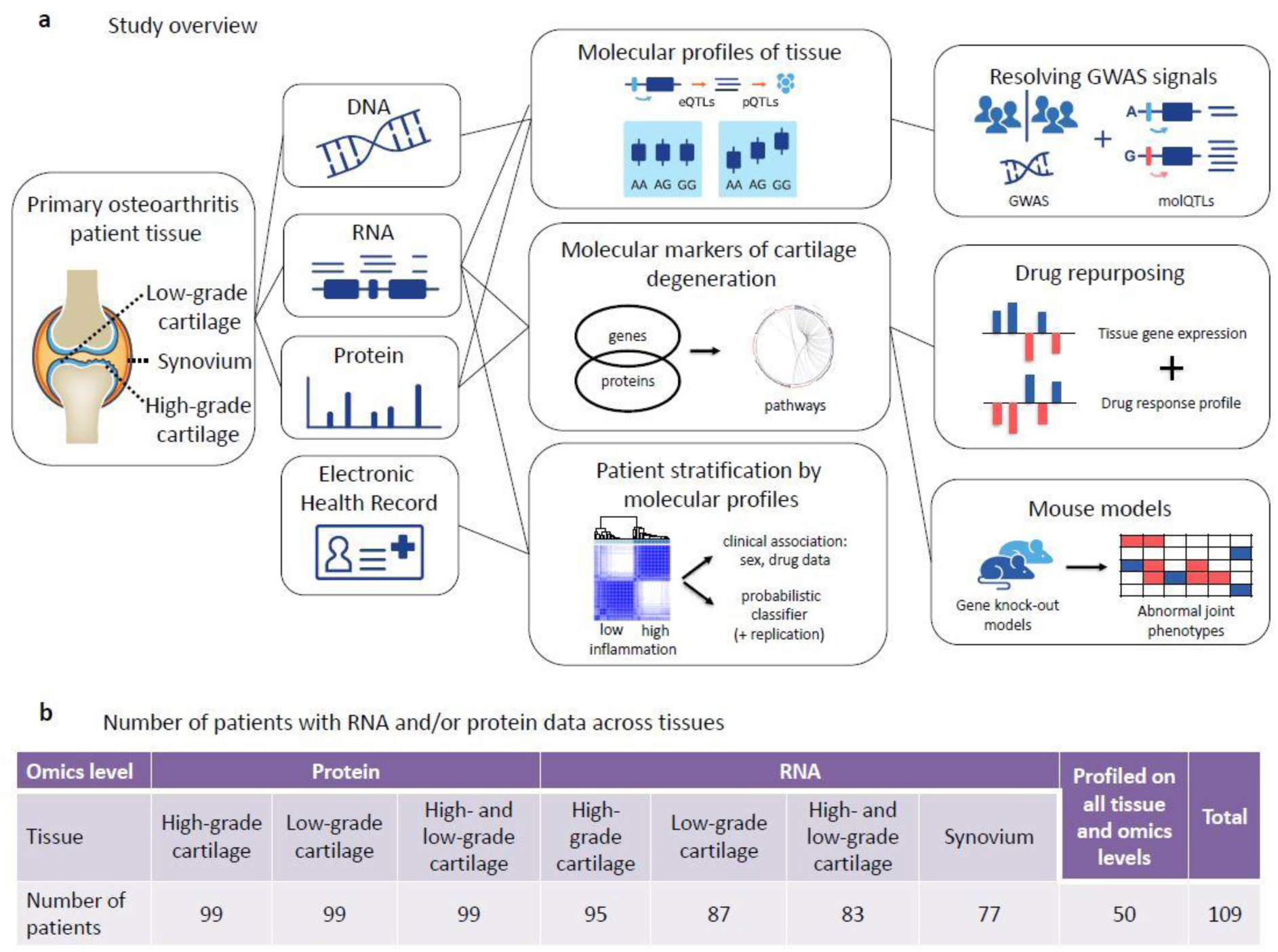
**Large-scale multi-omics characterisation of osteoarthritis disease tissue: study approach** a) We examined the molecular characteristics of osteoarthritis by profiling mRNA and proteins from low-grade cartilage, high-grade cartilage, and synovium tissue of over 100 patients undergoing total-joint-replacement for osteoarthritis, and combining these data with patient genotypes and information from electronic health records (EHRs). We identified genetic variants influencing mRNA or protein levels, several of which co-localise with genetic risk variants for osteoarthritis. We also identified molecular markers of cartilage degeneration, creating a gene expression profile of degeneration, and shortlisting existing drugs or compounds that reverse this profile in cell experiments. We generated mouse lines of several markers of cartilage degeneration, extensively profiling the bone and cartilage phenotypes of the mutant mice. Finally, we identified patient heterogeneity based on the molecular data, constructed and replicated a 7-gene probabilistic classifier to capture the heterogeneity, and identified associations with the patients’ clinical characteristics extracted from EHRs. b) Number of patients with data for each tissue and omics type after quality control. All 109 patients also have genome-wide genotype data.

Identification of molecular QTLs can help elucidate effector genes for genetic association signals, and provide a better understanding of the transcriptional regulation of key cell types in health and disease. For each gene, we considered genetic variants within 1Mb of the transcription start site, and followed a similar analysis approach to GTEx^6, 7^ (Methods). We identified *cis* expression QTLs (*cis*-eQTLs) for 1,891 genes in at least one tissue significant at 5% Storey-Tibshirani q-value^8^, with high correlation of effects across the tissues studied (Supplementary Figure 1a-c). The direction of effect was concordant across all *cis-*eQTLs detected in both low- and high-grade cartilage (92,758 variant-gene pairs: Pearson *r*=0.98, *P*<2.2x10^-16^). We identified *cis* protein QTLs (*cis*-pQTLs) for 38 genes in at least one tissue, with a similarly strong correlation across low- and high-grade cartilage (Pearson *r*=0.99, *P*<2.2x10^-16^, Supplementary Figure 1d-e, Supplementary Note).

To further identify differential regulation of gene expression between high- and low-grade cartilage, we examined variants with strong evidence for an eQTL effect in one tissue grade (posterior probability >0.9), but not in the other (posterior probability <0.1). We found 172 variants with differential effects on gene expression for 32 genes (differential eQTLs; Supplementary Table 1, Figure 2a-b). Sixteen genes had differential eQTLs located in a *cis*-acting regulatory region. Key genes in which this effect was observed were involved in development (transcription factor *HOXB2*), inflammation (*IL4I1*), and fibrosis (*CRLF1*). These genotype-dependent, divergent patterns of gene regulation between high- and low-grade cartilage underline the biological specificity of cell type and disease stage when investigating regulatory variant function.

**Figure 2.**
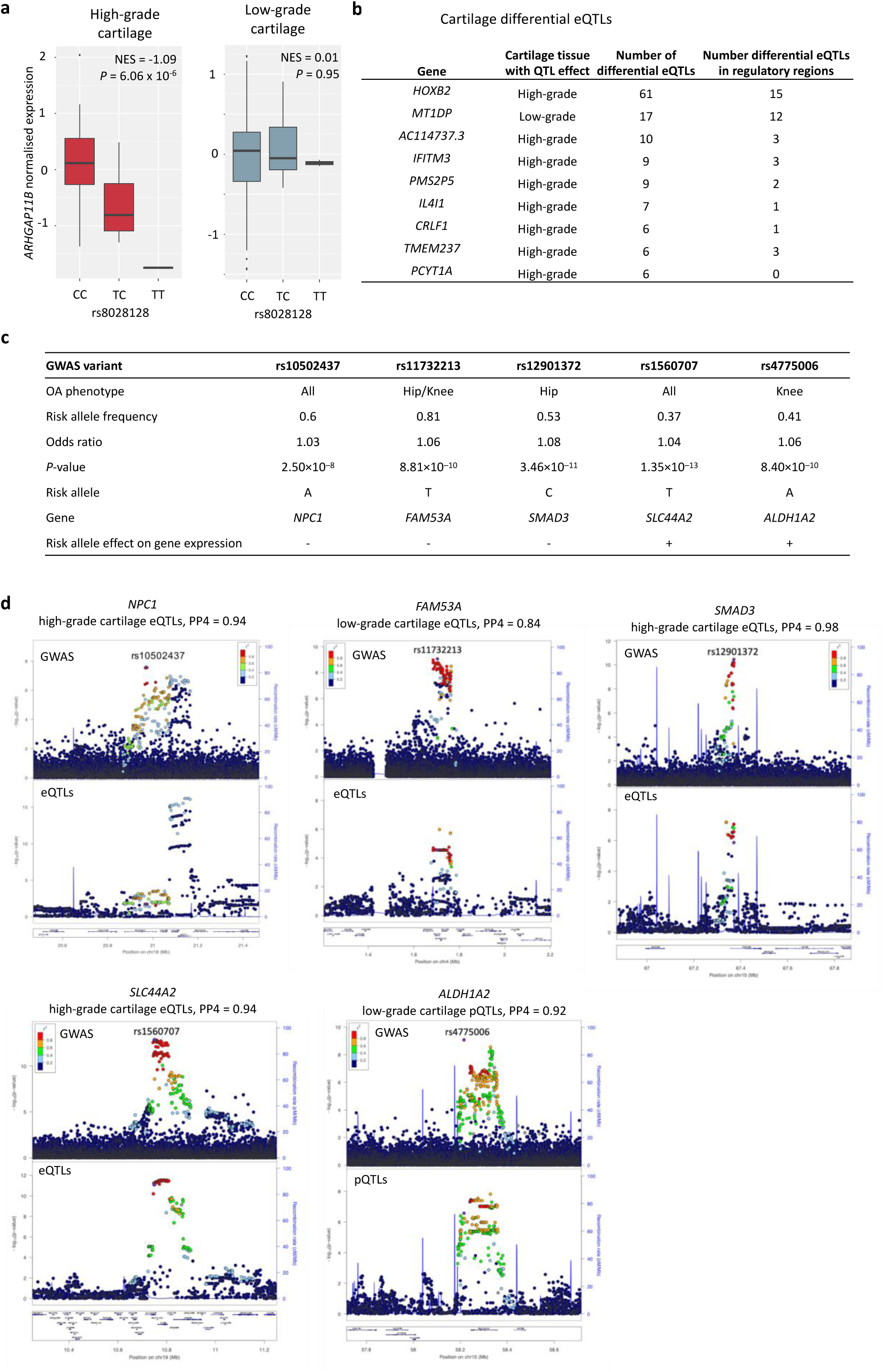
**Molecular QTLs in osteoarthritis disease tissue** a) An example of differential QTL effect: an association between genotype and gene expression present in high-grade (posterior probability m>0.9), but not low-grade cartilage (posterior probability m<0.1), or vice versa. Here, the association is present in high-grade, but not low-grade cartilage. The boxplots show normalised gene expression at 25^th^, 50^th^ and 75^th^ percentiles, and whiskers extend to 1.5 times the interquartile range. Inset: normalised effect size (NES) and FastQTL association *P*-value (*P*). b) Genes with at least 5 differential eQTL variants. Full results see Supplementary Table 1. c) Osteoarthritis GWAS signals with high posterior probability (≥0.80) for colocalisation with molecular QTLs. Each GWAS signal is denoted by its index variant. Risk allele effect: “+” for increase of expression with risk allele, “-“ for decrease. OA: osteoarthritis; AF: frequency of the risk allele; OR: odds ratio. d) GWAS and molecular QTL *P*-values in regions with colocalisation of the associations. Plots show 1Mb regions centered around the GWAS index SNPs (purple), with one point per genetic variant. PP4: posterior probability for colocalisation. For *NPC1* and *SMAD3*, colocalisation was also observed with low-grade cartilage molecular QTLs, with plots shown in Supplementary Figure 1f.

### Resolving GWAS signals

The majority of osteoarthritis genetic risk variants reside in non-coding sequence, making it challenging to identify the gene through which they confer their effect. Colocalisation analysis using molecular QTLs (molQTLs) can help clarify the mechanisms driving a GWAS locus by indicating whether the same variant is causal for both association with disease and for association with gene expression levels. We found strong evidence for colocalisation of 5 osteoarthritis loci with cartilage molQTLs for *ALDH1A2*, *NPC1*, *SMAD3*, *FAM53A*, and *SLC44A2* (posterior probability 0.84-0.98, Figure 2c-d). In all five instances, the GWAS index variant is non-coding. In three cases (*ALDH1A2, SMAD3* and *SLC44A2*) the likely effector gene is that residing closest to the lead variant. For the *NCP1* and *FAM53A* loci, the lead variants reside in introns of the *TMEM241* and *SLBP* genes, 141 kb and 18 kb away from the likely effector gene, respectively. This work helps pinpoint the identity of causal genes for hitherto unsolved association signals. In addition, our findings demonstrate the value of studying the relevant tissue and relevant stage for the disease under investigation. For example, when using data from GTEx resource^7^, which does not include cartilage, several osteoarthritis GWAS signals were found to co-localise with eQTLs in only 1 or 2 of 44 tissues (for example, *ALDH1A2*: ovary and tibial artery, *SMAD3*: skeletal muscle; *SLC44A2*: adrenal gland), without clear transferability of results to disease-relevant tissue. We have found robust evidence that that these three GWAS signals co-localise with molecular QTLs in cartilage.

To identify molecular signatures associated with disease severity, and hence likely effector genes for genetic association signals, we tested paired samples of high-versus low-grade cartilage for differential gene expression and protein abundance across 83 and 99 patients, respectively. We detected significant gene expression differences for 2,557 genes at 5% false discovery rate (FDR; Figure 3a), and protein abundance differences for 2,233 proteins at 5% FDR (Figure 3a, Supplementary Figure 2, Methods, Supplementary Note). 409 of these genes (Supplementary Table 2) demonstrated significant differential expression at both the RNA and protein levels, lending robust, cross-omics evidence for involvement in disease progression. We found strong evidence for concordant direction of expression changes across the two omics levels, with a correlation of *r*=0.63 (*P*<1.0x10^-17^) between the RNA- and protein-level effect sizes for genes with cross-omics changes (Figure 3b). In keeping with previous, smaller-scale reports^9–11^, extracellular matrix (ECM)-receptor interaction emerged as the primarily activated pathway in high-compared to low-grade cartilage (Methods, Figure 3c, Supplementary Note, Supplementary Table 3, Supplementary Figure 2).

**Figure 3.**
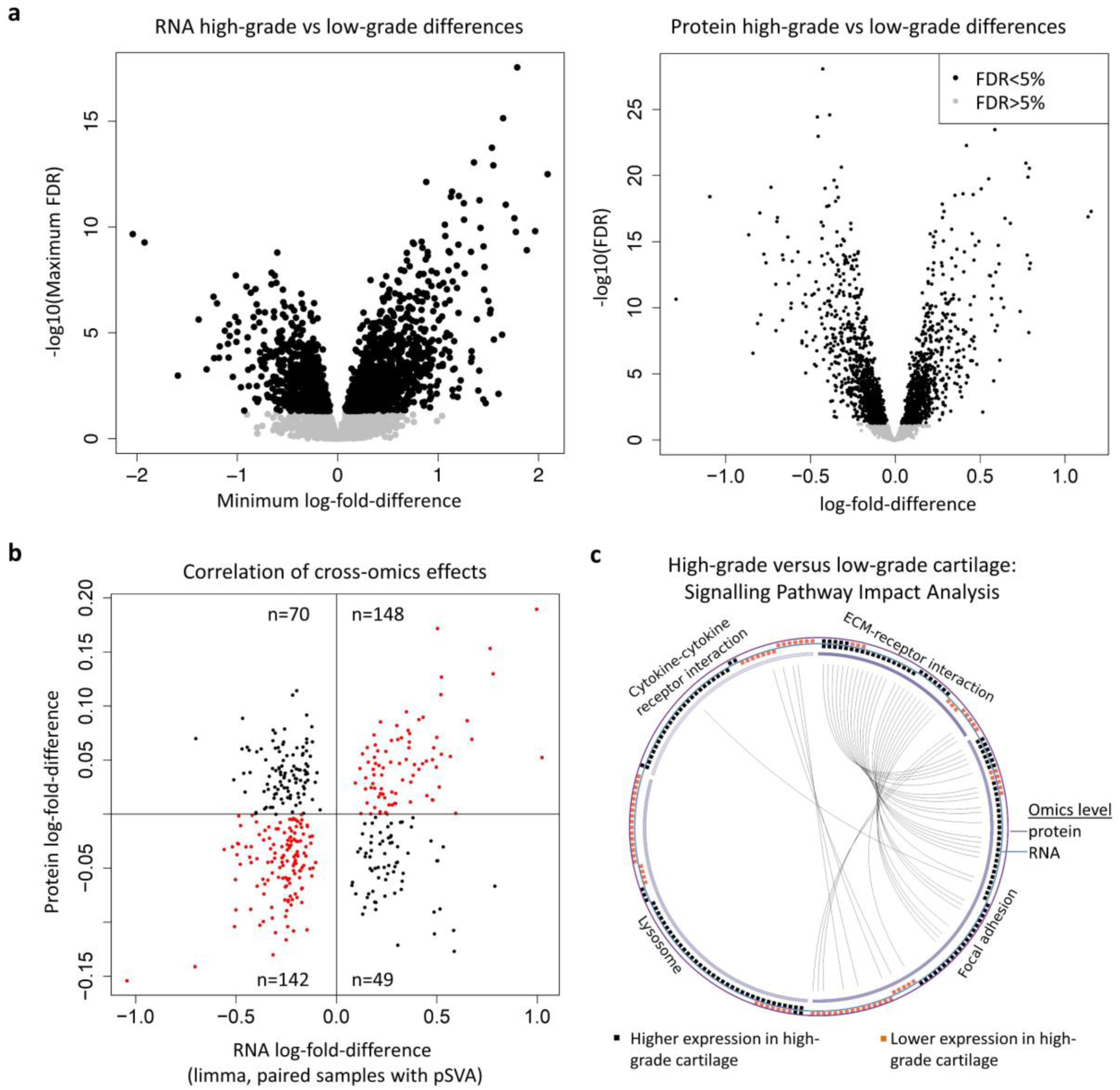
**Molecular differences between high-grade and low-grade cartilage** a) Wide-spread RNA-level (left) and protein-level (right) differences between high-grade and low-grade cartilage. The RNA plot shows conservative results based on different approaches (see Methods), with 2,557 differentially expressed genes significant at 5% FDR in all approaches. b) RNA- and protein-level log-fold differences for 409 genes with significant cross-omics differences between high-grade and low-grade cartilage (see Supplementary Figure 2a for all genes). The direction of difference agrees for 290 of the 409 genes (71%; binomial *P*<1.0x10^-17^), with a strong correlation of effect sizes (Pearson *r*=0.63, *P*<10^-10^). c) Signalling Pathway Impact Analysis (SPIA) identified biological pathways associated with differences between high-grade and low-grade cartilage. Pathways with significant results at 5% FDR based on RNA-level changes are shown, all activated in high-grade cartilage. Boxes on the outside circles represent individual genes, with arches connecting the same gene across pathways. See also Supplementary Figure 2b and full results in Supplementary Table 3.

We found 91 of the genes with significantly different expression profiles between high- and low-grade cartilage to also be associated with genetic risk of osteoarthritis^5^ (gene-level *P*-value significant after Bonferroni correction for multiple testing at *P*<1.02x10^-6^; Methods, Supplementary Table 4). For example, variants in *ALDH1A2* are associated with knee osteoarthritis, and we found significantly higher *ALDH1A2* gene expression and lower protein expression levels in high-grade cartilage. For *SLC39A8*, the GWAS signal was fine-mapped to a single missense variant with posterior probability of 0.999 and the gene demonstrated higher expression levels in high-grade cartilage. These findings highlight the value of integrating multi-omics data with genetic association summary statistics to identify likely effector genes for GWAS signals.

### Patient stratification

Better stratification of patients by molecular endotype can provide opportunities for tailored therapeutic intervention. The analysis of primary tissue samples offers the possibility to stratify patient groups on the basis of their molecular profiles. Previous studies in smaller patient sets have identified discrete subgroups by using gene expression arrays or RNA sequencing in low-grade cartilage^12, 13^. Here, we substantially increased sample size and hence power, and were able to test for overlap between clusters identified from different tissues, assess the impact of cluster assignment on differences between high- and low-grade cartilage, and evaluate whether the clustering is truly categorical or better represented by a continuous spectrum of variation.

First, we applied a consensus clustering analysis to identify discrete subgroups across patient tissue samples. Based on RNA sequencing data, we identified 2 robust patient clusters in synovium (42 and 34 individuals, respectively), each of which further formed 2 sub-clusters (Figure 4a, Supplementary Figure 3a,c). We identified 2 robust patient clusters within low-grade cartilage (45 and 42 individuals, respectively; Figure 4b, Supplementary Figure 3a,c), and no clear sub-clustering within high-grade cartilage (Supplementary Figure 3a,b). Cartilage clustering was independent of the synovium clusters (Fishers *P*-value >0.66).

**Figure 4.**
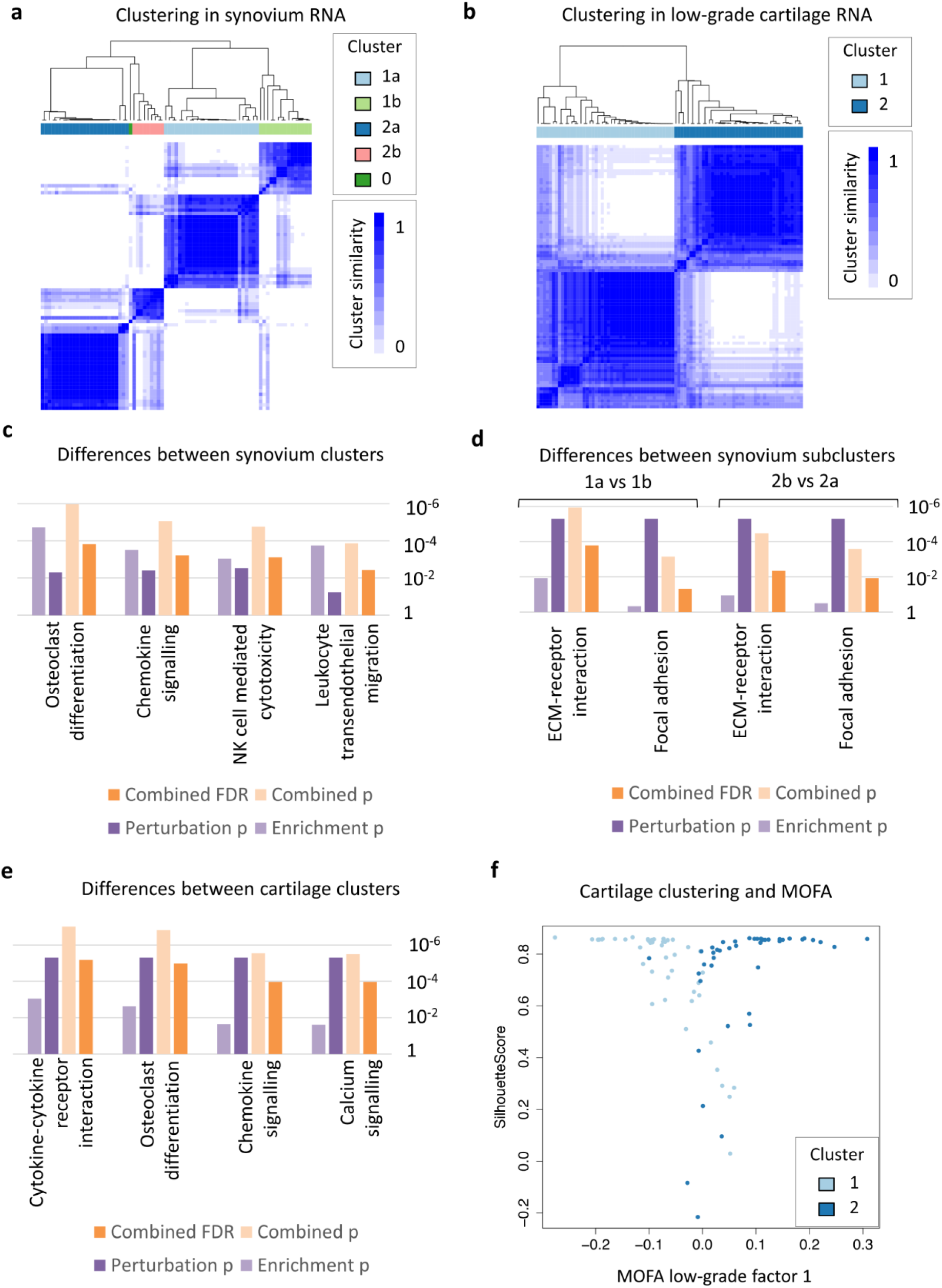
**Distinct clusters identified in low-grade cartilage and synovium tissue** a) Synovium tissue samples from patients are separated into two clusters based on consensus clustering of RNA data (synovium-Cluster1, *n*=42 and synovium-Cluster2, *n*=34). Each cluster formed 2 sub-clusters (synovium-Cluster1a, *n*=27 and synovium-Cluster1b, *n*=15; separately synovium-Cluster2a, *n*=25 and synovium-Cluster2b, *n*=9). Cluster 0: one outlier sample. b) Low-grade cartilage tissue samples from patients are separated into two clusters based on consensus clustering of RNA data (cartilage-Cluster1, *n*=45 and cartilage-Cluster2, *n*=42). c) Gene expression differences between synovium clusters show several significant (FDR<5%) associations related to inflammation and osteoclast differentiation using Signalling Impact Pathway Analysis (SPIA). d) Gene expression differences between the synovium sub-clusters within each cluster show similar pathway associations, including to ECM-receptor interaction and focal adhesion pathways. e) Gene expression differences between low-grade cartilage clusters show several significant pathway associations, including inflammation and osteoclast differentiation. f) An analysis of low-grade cartilage samples using MOFA identifies a continuous spectrum of variation between samples. Samples with high MOFA Factor 1 scores are mostly in cartilage-Cluster1 and those with low MOFA Factor 1 scores mostly in cartilage-Cluster2. Samples with intermediate MOFA Factor 1 scores have lower Silhouette Scores, showing more uncertainty in cluster assignment. For synovium, see Supplementary Figure 4c. For c)-e), the barplots show SPIA *P*-values and FDRs for for over-representation analysis of genes (“Enrichment p”); perturbation of the pathway based on gene log-fold differences (“Perturbation p”); combining enrichment and perturbation *P*-values (“Combined p” and “Combined FDR”). The associations shown are robust across several gene-level differential expression cut-offs (Supplementary Table 5).

Gene expression analysis showed large differences between the synovium clusters and sub-clusters, with over 5,000 genes differentially expressed at 5% FDR (Supplementary Figure 4). Signalling pathway impact analysis showed that the differences between the two patient clusters relate to inflammation, while differences between the sub-clusters relate to the extracellular matrix and to cell adhesion (with pathway associations significant at 5% FDR, see Methods, Figure 4c-d, Supplementary Table 5). Gene expression analysis also identified strong differences between the two low-grade cartilage clusters, with over 7,500 genes differentially expressed at 5% FDR. This clustering was also strongly associated with inflammation, extracellular matrix-related and cell adhesion pathways (FDR<5%; Figure 4e, Supplementary Table 5). Our data lend robustness to evidence^12, 13^ that inflammation plays a role in ostearthritis subtype identification, and we show that this finding emerges from both cartilage and synovial tissue analysis (Supplementary Note). The presence of an inflammatory endotype axis within osteoarthritis raises the possibility for patient selection to clinical trials of inflammation-modulating investigational therapies in appropriately identified patients.

In the within-cluster comparison of high-grade and low-grade cartilage samples, the gene expression differences showed high correlation with unstratified analysis of all patients (log-fold differences Pearson *r*=0.89, Spearman *ρ*=0.89, *P*<10^-10^ for cartilage-Cluster 1; and Pearson *r*=0.93, Spearman *ρ*=0.92, *P*<10^-10^ for cartilage-Cluster 2). Conversely, among genes differentially expressed in the all-patient analysis, 99% had the same direction of effect in the analysis of patients within each low-grade cartilage cluster. These data indicate that gene expression differences between high- and low-grade cartilage are independent of low-grade cartilage cluster (Supplementary Figure 4a).

Disease endpoints and discrete endotypes represent underlying processes that may be more sensitively captured by continuous axes of variation within the molecular data rather than categorical classifiers. Such an approach may help define disease trajectories earlier in the natural history of osteoarthritis. To evaluate this, we applied multi-omics factor analysis (MOFA)^14^, an integrative method that can discover hidden factors that represent drivers of variability between samples or patients (latent factors) that is akin to a cross-data principal component analysis. Cross-omics analysis of patients showed that the first two factors (axes of variation) were strongly associated with immune system processes and the extracellular matrix (Methods, Supplementary Note), in keeping with the biological pathways identified to play an important role above. We also found the continuous axes of variation within low-grade cartilage and synovium to correspond strongly with cluster assignment (Figure 4f, Supplementary Figures 4c,5, Supplementary Note). This is consistent with variation within tissues being better captured as a continuous spectrum rather than as discrete clusters.

### Molecular classifier

To allow characterisation of future patients within the inflammatory endotype axis for potential personalised treatment, we sought to develop a molecular tool that can predict patient assignment based on a small number of genes. To this end, we applied a soft-thresholding centroid-based method, PAMR^15^, and identified 7 genes, the expression levels of which could be combined to predict cluster assignment for each patient sample (Figure 5a, Supplementary Figure 6): *MMP1*, *MMP2*, and *MMP13*, known to be involved in cartilage degradation^16^; *IL6*, a pro-inflammatory cytokine; *CYTL1*, a cytokine-like gene, loss of which has been found to augment cartilage destruction in surgical osteoarthritis mouse models^17^; *APOD*, a component of high-density lipoprotein found to be strongly up-regulated by retinoic acid^18^, which is in turn regulated by *ALDH1A2*^19^, an osteoarthritis risk locus^5, 20^; and *C15orf48*, of currently unknown function. Notably, the posterior probabilities for cluster assignment generated by the classifier captured the main continuous spectrum of variation in this tissue (Spearman correlation *ρ*=-0.95, *P*<10^-10^; Figure 5b).

**Figure 5.**
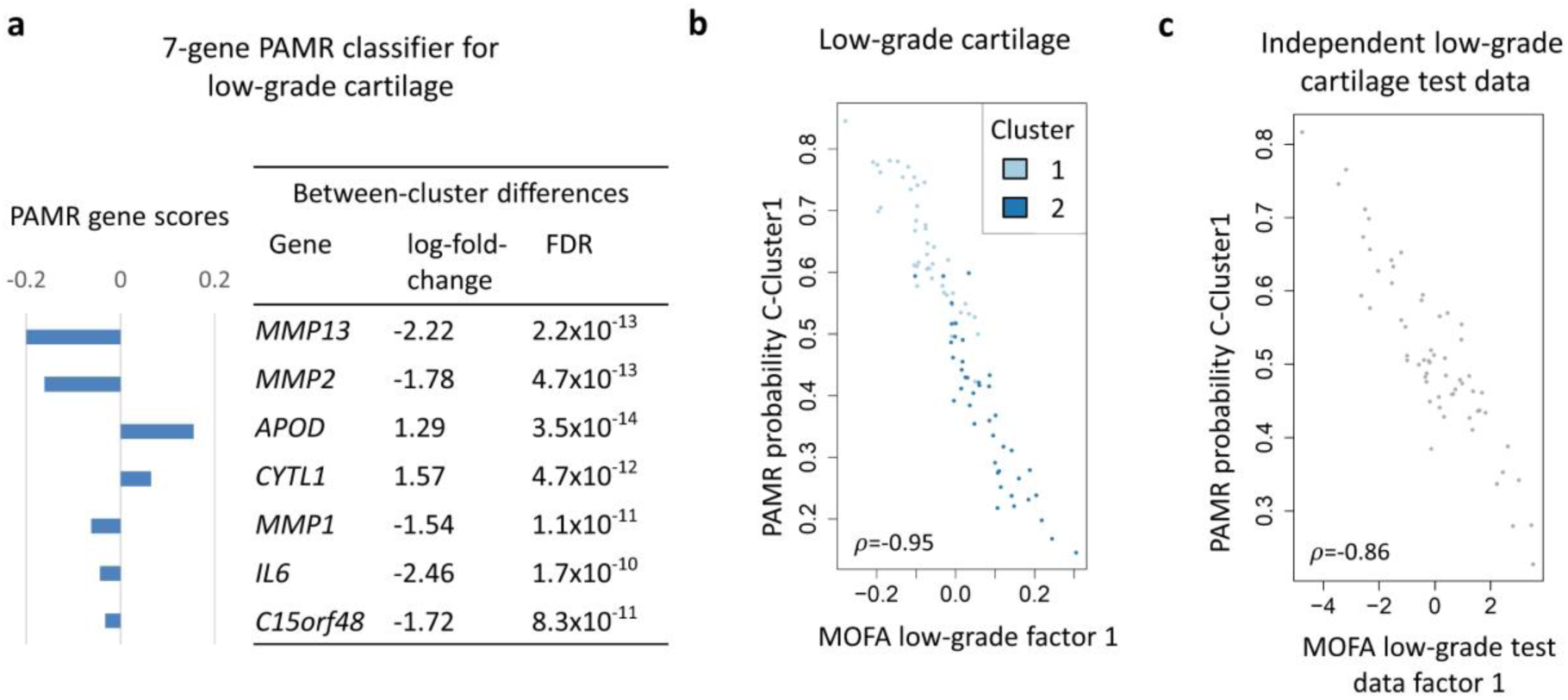
**Variation within low-grade cartilage can be recovered using a 7-gene classifier** a) Using PAMR, we constructed a 7-gene classifier to predict cluster assignment for low-grade cartilage samples. The barplot shows the PAMR score for each gene (the difference between the standardised centroids of the two clusters), the right panel the differential expression of the genes between the two low-grade cartilage clusters. See also Supplementary Figure 6 for classifier performance. b) The PAMR posterior probabilities for cluster assignment are highly correlated with MOFA Factor 1 scores for low-grade cartilage samples, capturing the continuous spectrum of variation between samples. Inset: Spearman correlation, *P*<10^-10^. c) In an independent set of 60 low-grade cartilage samples from 60 osteoarthritis patients undergoing total-knee-replacement, the posterior probabilities for cluster assignment from the 7-gene classifier are well-correlated with the continuous spectrum of variation in these samples, as quantified by MOFA Factor 1 in an *ab initio* analysis. Inset: Spearman correlation, *P*<10^-10^.

We validated the 7-gene classifier in an independent gene expression dataset of low-grade cartilage samples from 60 knee osteoarthritis patients undergoing joint replacement surgery, which had also identified separation of samples into two groups^13^. This group assignment corresponded to the 7-gene classifier categorical cluster assignment for 73% of the samples (32 out of 44 samples with available data). The posterior probabilities for cluster assignment had good correspondence to the main continuous spectrum of variation (Spearman correlation *ρ*=-0.86, *P*<10^-10^; Figure 5c). Of the genes used to distinguish the two groups in the other study, the majority showed a discordant direction of effect or higher false-discovery rates (33.5-99.8% FDR) between the low-grade cartilage clusters in this work (Supplementary Note, Supplementary Table 6). By contrast, all 7 genes used by the 7-gene classifier showed concordant differences between the two groups from the other study (*MMP1*, *MMP2*, *MMP12* showed significant differences at 0.07% FDR; *APOD*, *CYTL1*, *C15orf48* at 11-15% FDR; *IL6* at 25.5% FDR), indicating improvement in transferrability. These findings support the predictive potential of the 7-gene classifier.

### Clinical profiles of molecular clusters

We investigated whether the stratification of patients into different tissue-based transcriptional profile clusters was associated with clinical characteristics. We compiled information on sex, age, height, weight, body mass index, and electronic health records information on pre-operative American Society of Anesthesiologists (ASA) grade^21^ and prescribed medications at the time of joint replacement surgery. Cartilage cluster assignment was associated with patient sex (OR=4.12, *P*=2.4x10^-3^), with women more likely to be members of the cluster characterised by higher inflammation. One explanation for this observation may be the lower concentration of oestrogen and androgens, which have established anti-inflammatory effects, in post-menopausal women^22–24^. This is also in line with the disproportionate increase in the incidence of osteoarthritis in women after the menopause. Patients in the high-inflammation cluster were also more likely to be prescribed proton pump inhibitors (OR=4.21, *P*=4.0x10^-3^; Supplementary Table 7). Several further clinical characteristics were associated with cluster assignment at nominal significance: patients in the high-inflammation cluster were more likely to be prescribed a higher number of drugs (OR=1.21 per additional drug, *P*=0.023) and to be older (OR=1.06 per year, *P*=0.036). To verify the robustness of these associations, we carried out a sensitivity analysis explicitly accounting for sex or sex and age, and observed qualitatively and quantitatively similar effects (Methods, Supplementary Table 7). These findings support a mechanistic explanation of the established association between osteoarthritis, age, sex, multimorbidity and polypharmacy^25, 26^, although the direction of causation remains unclear.

### Candidate therapeutic compounds

We aimed to identify compounds with the potential to reverse the spectrum of molecular differences between high- and low-grade patient cartilage based on existing *in vitro* drug screen data. We used ConnectivityMap^27^, a dataset of 2,684 gene expression perturbations induced by compounds across 9 human cell lines, to assess each perturbation profile against our differentially expressed genes using the clue.io platform. We identified 19 compounds that induced strong opposing gene expression signatures to the differences between high- and low-grade cartilage, reducing the expression of genes with cross-omics higher expression in high-grade cartilage (summary tau and median tau below -0.95, see Methods, Table 1, Supplementary Table 8). These identified compounds include oestrogen receptor agonists diethylstilbestrol and alpha-estradiol, the latter of which targets *KCNMA1*, coding for the pore-forming alpha subunit of a calcium-sensitive potassium channel that demonstrated significantly lower gene expression and protein abundance in high-grade cartilage. These findings are consistent with our clinical classifier, molecular clustering, and with established epidemiological data showing an association between osteoarthritis and oestrogen deficiency^28^. Although studies of oestrogen therapy for osteoarthritis have been inconclusive^29, 30^, identification of cartilage-specific oestrogen-mediated pathways, such as through *KCNMA1*, may allow more focussed investigational molecule development.

**Table 1.** Compounds with strongest evidence for inducing gene expression signatures that counter differences between high-grade and low-grade cartilage. Results based on data from ConnectivityMap^27^ and the clue.io platform. DE targets: drug targets as listed in ConnectivityMap with differential expression between high-grade and low-grade cartilage on RNA (R) or protein (P) level, “+” and “-“ indicate higher or lower expression high-grade cartilage, respectively. The 10 compounds with lowest median tau scores are shown; the full list of compounds is in Supplementary Table 8.

| Name | Description | DE targets |
| --- | --- | --- |
| Emetine | protein synthesis inhibitor | RPS2 (P+) |
| Rucaparib | PARP inhibitor | PARP2 (R-) |
| Alpha-estradiol | estrogen receptor agonist | KCNMA1 (R-, P-) |
| VEGF-receptor-2-kinase-inhibitor-IV | VEGFR inhibitor |  |
| IB-MECA | adenosine receptor agonist,<br>granulocyte colony stimulating<br>factor agonist |  |
| Diethylstilbestrol | estrogen receptor agonist, chloride<br>channel blocker |  |
| KIN001-220 | Aurora kinase inhibitor |  |
| SB-216763 | glycogen synthase kinase inhibitor | GSK3B (R+, P+), CDK2<br>(P-) |
| RHO-kinase-inhibitor-III[rockout] | ROCK inhibitor | IMPDH2 (P-) |
| Nornicotine | acetylcholine receptor agonist |  |

Several further compounds with known biological relevance to osteoarthritis were also identified by the ConnectivityMap analysis (Table 1): IB-MECA (an adenosine receptor agonist used as an anti-inflammatory drug in rheumatoid arthritis) (https://pubchem.ncbi.nlm.nih.gov/compound/ib-meca), VEGF-receptor-2-kinase-inhibitor-IV, RHO-kinase-inhibitor-III[rockout] (a rho associated kinase inhibitor), and nornicotine (an acetylcholine receptor agonist extracted from tobacco and related to nicotine) (https://pubchem.ncbi.nlm.nih.gov/compound/nornicotine). In a rat model of chemically-induced osteoarthritis, IB-MECA prevented cartilage damage, osteoclast/osteophyte formation, and bone destruction^31^. *VEGF* modulates chondrocyte survival during development and is essential for bone formation and skeletal growth. However, dysregulation of *VEGF* expression in the adult joint is a feature of osteoarthritis^32^. Conditional knock-down of *Vegf* attenuates surgically-induced osteoarthritis in mice, with intra-articular anti-VEGF antibodies as well as oral administration of the VEGFR2 kinase inhibitor Vandetanib suppressing osteoarthritis progression^33^. In a rat model of osteoarthritis, a rho kinase inhibitor was found to reduce knee cartilage damage^34^. Finally, there is a well-established effect of smoking on osteoarthritis^5, 35^. Together, these results identify candidate compounds that warrant investigation, and provide evidence for the validity of this approach.

In addition to signatures induced by compounds, ConnectivityMap contains gene expression profiles induced by *in vitro* gene knock-down or over-expression. We identified 36 genes for which the experimental perturbation induces changes in the opposite direction to molecular differences between high- and low-grade cartilage (Supplementary Table 8), notably including knock-down of *IL11*. Variation in *IL11* is associated with increased risk of hip osteoarthritis^5^, and the gene is up-regulated in osteoarthritis knee tissue^36^, with a similar trend observed here. IL11 is a cytokine with a key role in inflammation, and monoclonal anti-IL11 antibodies have been developed for use in several diseases. These findings provide strong supportive evidence for down-regulation of *IL11* as a potential therapeutic intervention for osteoarthritis.

Better understanding of the molecular landscape of osteoarthritis has provided the basis for shortlisting drugs whose expression signatures show the potential to reverse the spectrum of molecular differences between high- and low-grade cartilage. This screening approach could facilitate further refinement of existing compound groups to enhance their biological activity within osteoarthritis pathways where existing agents are of uncertain efficacy.

## Discussion

Osteoarthritis is a globally important condition of huge public health relevance. As a heterogeneous disease, it requires patient stratification for successful therapy development and translation. Here, we have combined genome-wide genotyping with RNA sequencing and quantitative proteomics in primary human tissues to construct the first deep molecular quantitative trait locus map of cell types directly involved in the disease. By integrating multiple layers of omics data, we have helped resolve genetic association signals and identified likely effector genes.

Our findings highlight the importance of generating molQTL data in disease-relevant tissue, and add to the growing body of evidence that disease risk variants can act over long distances, including within introns of one gene while influencing the expression of another gene. The availability of molecular QTL data offers a resource that will enable the resolution of further genetic association signals emerging from ongoing large-scale efforts in osteoarthritis (https://www.genetics-osteoarthritis.com/) and further diseases and traits of musculoskeletal relevance, in which chondrocytes and synoviocytes are cell types of importance.

In this work, we have further identified key biological processes driven by molecular differences between high- and low-grade cartilage based on differentially expressed genes. For 7 of the genes with significant expression differences, we generated genetically-modified mice with mutant alleles in orthologues and carried out detailed joint phenotyping of adult mice from these 7 lines^37^ (Methods). We identified at least one abnormal joint phenotype at nominal significance (*P*<0.05) for each of the 7 genes studied (Supplementary Figures 7-8, Supplementary Note), functionally validating their role in joint morphology.

A key consideration in the selection of patients for clinical trials is the appropriate targeting of individuals at highest risk of disease progression, and how these patients can be identified. Several clinical risk factors for progression (such as age, ethnicity, BMI, infrapatellar fat pad synovitis, and co-morbidity count) have been well-described^38^. The association of molecularly-defined patient clusters with some of these clinical characteristics lends evidence to support the integration of genomic biomarkers to drive precision medicine approaches in osteoarthritis, and suggests a molecular basis for targeting disease-modifying interventions. Our analysis of patient clustering on the basis of chondrocyte gene expression profiles is consistent with previous work showing that individuals may be clustered by inflammation. Here, by virtue of the added power of a substantially larger sample size, we show that variation within the tissues is better captured by a continuous spectrum, rather than as discrete clusters. The 7-gene classifier constructed in this work and validated using independent data can place patients along the inflammatory endotype axis of variation. Such molecular endotyping approaches may thus have the potential to be applied for targeting treatment to the right patients and at the right time, although further study will first be required to determine to which extent the inflammation axis is stable across time or differs with disease activity.

The molecular stratification of these patient clusters would require validation of the the 7-gene classifier tool for tissue that can be non-destructively collected and that is acceptable to patients, such as saliva, blood, or synovial fluid. Going forward, establishing the classifier’s ability to identify differential rates of clinical disease progression in longitudinal studies referenced against a robust clinical or radiological disease progression endpoint would be warranted to investigate potential utility as a clinical tool.

In summary, by integrating multiple layers of omics data, we have helped resolve genetic association signals by identifying likely effector genes, and have highlighted opportunities for new high-value targets. We demonstrate how integrating multi-omics in primary human complex disease tissue can serve as a valuable approach that moves from basic discovery to accelerated translational opportunities. Our findings identify drug repurposing opportunities and allow the identification of novel investigational avenues for patient stratification, disease severity, and therapy development, responding to the global clinical challenge of osteoarthritis.

## Acknowledgements

We thank the study participants who made this work possible by their generous donation of samples. This work was funded by the Wellcome Trust (206194). M.J.C. was funded through a Centre for Integrated Research into Musculoskeletal Ageing grant (MRC 148985). R.A.B. and the Human Research Tissue Bank are supported by the NIHR Cambridge Biomedical Research Centre. J.H.D.B. and G.R.W. are funded by a Wellcome Trust Strategic Award (101123), a Wellcome Trust Joint Investigator Award (110140 and 110141) and a European Commission Horizon 2020 Grant (666869, THYRAGE). A.W.M. receives funding from Versus Arthritis; Tissue Engineering and Regenerative Therapies Centre (21156). Mutant mice were generated via Wellcome Trust grant WT098051. We thank members of the Sanger Institute Mouse Pipelines teams (Mouse Informatics, Molecular Technologies, Genome Engineering Technologies, Mouse Production Team, Mouse Phenotyping) and the Research Support Facility for the provision and management of the mice. This research was conducted using the UK Biobank Resource under application number 9979. The authors are grateful to Dr. Iris Fischer for helpful edits.

## Author Contributions

Study design: E.Z., J.M.W, J.S. Collection of knee samples: M.J.C., R.L.J., D.S., K.S., J.M.W. Collection of hip samples: R.A.B., A.W.M. Review of patient electronic health record data: A.F., J.M.W. Proteomics assays: T.I.R., J.S.C. Mouse Resources: C.J.L. Mouse experiments: N.B., K.F.C., S.M.P., J.H.D.B., G.R.W. Molecular QTL and colocalisation analyses: L.S. Differential expression analyses: J.S., L.S. Pathway association, tissue clustering, MOFA, drug repurposing, and statistical mouse data analyses: J.S. Writing - original draft: J.S, L.S., J.M.W., E.Z. Writing - comments and review: all authors.

## Online Methods

### Data reporting

No statistical methods were used to predetermine sample size. The experiments were not randomized and the investigators were not blinded to allocation during experiments and outcome assessment.

### Study participants

We collected tissue samples from 115 patients undergoing total joint replacement surgery in 4 cohorts: 12 knee osteoarthritis patients (cohort 1; 2 women, 10 men, age 50-88 years, mean 68 years); 20 knee osteoarthritis patients (cohort 2; 14 women, 6 men, age 54-82 years, mean 70 years); 13 hip osteoarthritis patients (cohort 3; 8 women, 5 men, age 44-84 years, mean 62 years); 70 knee osteoarthritis patients (cohort 4; 42 women, 28 men, age 38-84 years, mean 70 years).

All patients provided written, informed consent prior to participation in the study. Matched low-grade and high-grade cartilage samples were collected from each patient, while synovial lining samples were collected from patients in cohorts 2 and 4.

#### Cohorts 1, 2, 4 (knee osteoarthritis)

This work was approved by Oxford NHS REC C (10/H0606/20 and 15/SC/0132), and samples were collected under Human Tissue Authority license 12182, Sheffield Musculoskeletal Biobank, University of Sheffield, UK.

We confirmed a joint replacement for osteoarthritis, with no history of significant knee surgery (apart from meniscectomy), knee infection, or fracture, and no malignancy within the previous 5 years. We further confirmed that no patient used glucocorticoid use (systemic or intra-articular) within the previous 6 months, or any other drug associated with immune modulation. For cohort 1, cartilage samples were scored using the OARSI cartilage classification system9,10. From each patient, we obtained one sample with high OARSI grade signifying high-grade degeneration (“high-grade sample”), and one cartilage sample with low OARSI grade signifying healthy tissue or low-grade degeneration (“low-grade sample”).

For cohorts 2 and 4, cartilage samples were scored macroscopically using the International Cartilage Repair Society (ICRS) scoring system11. From each patient, we obtained one sample of ICRS grade 3 or 4 signifying high-grade degeneration (“high-grade sample”), and one cartilage sample of ICRS grade 0 or 1 signifying healthy tissue or low-grade degeneration (“low-grade sample”). For cohorts 2 and 4, we also collected synovial membrane from the suprapatellar region of the knee joint.

Finally, from all patients in cohorts 1,2, and 4, we also obtained a blood sample to extract DNA for genotyping.

We obtained information on patient clinical characteristics (age, height, weight, body mass index (BMI), American Society of Anaesthesiologists (ASA) grade^21^) from the electronic patient records. For each patient, a list of drugs prescribed on the date of sample collection was also compiled from the electronic patient record and cross referenced with the patient medical history.

#### Cohort 3 (hip osteoarthritis)

Samples were collected under National Research Ethics approval reference 11/EE/0011, Cambridge Biomedical Research Centre Human Research Tissue Bank, Cambridge University Hospitals, UK.

We confirmed osteoarthritis disease status by examination of the excised femoral head. From each patient, we obtained a cartilage sample showing a fibrillated or fissured surface signifying high-grade degeneration (“high-grade sample”), one cartilage sample showing a smooth shiny appearance signifying healthy tissue or low-grade degeneration (“low-grade sample”).

### Isolation of chondrocytes

#### Cohorts 1,2,4

We followed a previously established protocol to isolate chondrocytes^9^. Osteochondral samples were transported in Dulbecco’s modified Eagle’s medium (DMEM)/F-12 (1:1) (Life Technologies) supplemented with 2mM glutamine (Life Technologies), 100 U/ml penicillin, 100 μg/ml streptomycin (Life Technologies), 2.5 μg/ml amphotericin B (Sigma-Aldrich) and 50 μg/ml ascorbic acid (Sigma-Aldrich) (serum free media). Half of each sample was then taken forward for chondrocyte extraction. Cartilage was removed from the bone, dissected and washed twice in 1xPBS. Tissue was digested in 3 mg/ml collagenase type I (Sigma-Aldrich) in serum free media overnight at 37°C on a flatbed shaker. The resulting cell suspension was passed through a 70 μm cell strainer (Fisher Scientific) and centrifuged at 400g for 10 minutes. Subsequently, the cell pellet was washed twice in serum free media and centrifuged at 400g for 10 minutes. The resulting cell pellet was resuspended in serum free media. Cells were counted using a haemocytometer and the viability checked using trypan blue exclusion (Invitrogen). The optimal cell number for spin column extraction from cells was between 4x10^6^ and 1x10^7^. Cells were then pelleted and homogenized.

#### Cohort 3

The extraction of chondrocytes in the majority of these samples has previously been described^39^, with the remaining samples following the same protocol. The protocol was based on that for cohorts 1,2,4 and highly similar. Briefly, each cartilage portion was minced with a scalpel and placed in 20ml of Dulbecco’s modified Eagle medium (Invitrogen) containing 10% foetal bovine serum (Invitrogen) and 6 mgml^-1^ collagenase A (Sigma). The tissue culture flasks were incubated overnight to digest the cartilage pieces. The resulting cell suspension was passed through a 30 μm filter (Miltenyi) and centrifuged at 400g for 10 minutes. The cell pellet was then re-suspended in 1ml of PBS and counted on a haemocytometer following 1:1 mixing with trypan blue to determine cell viability.

### Isolation of synoviocytes

We followed a previously established protocol to process synovial samples^40^. Synovial samples were transported in serum free media, as described above. The synovial membrane was dissected from underlying tissue then trypsinised for 1 hour. Tissue was then digested in 1mg/ml Collagenase Blend H (Sigma Aldrich) in serum free media overnight at 37°C on a flatbed shaker. The resulting cell suspension was passed through a 100 μm cell strainer (Fisher Scientific) and centrifuged at 400g for 10 minutes. Subsequently, the cell pellet was washed twice in serum free media and centrifuged at 400g for 10 minutes. The resulting cell pellet was resuspended in serum free media. Cells were counted using a haemocytometer and the viability checked using trypan blue exclusion (Invitrogen). The optimal cell number for spin column extraction from cells was between 4x10^6^ and 1x10^7^. Cells were then pelleted and homogenized.

### DNA, RNA and protein extraction

DNA, RNA, and protein extraction was carried out using Qiagen AllPrep DNA/RNA/Protein Mini Kit following manufacturer’s instructions, with small variations for cohort 3 as previously described^39^. Samples were frozen at -80°C (cohorts 1, 2, 4) or -70°C (cohort 3) prior to assays.

### RNA sequencing

We performed a gene expression analysis on samples from 113 patients (Supplementary Table 9). We purified poly-A tailed RNA (mRNA) from total RNA using Illumina’s TruSeq RNA Sample Prep v2 kits. We then fragmented the mRNA using metal ion-catalyzed hydrolysis and synthesized a random-primed cDNA library. The resulting double-strand cDNA was used as the input to a standard Illumina library prep, whereby ends were repaired to produce blunt ends by a combination of fill-in reactions and exonuclease activity. We performed A-tailing to allow samples to be pooled, by adding an “A” base to the blunt ends and ligation to Illumina Paired-end Sequencing adapters containing unique index sequences. Due to better performance, the 10-cycle PCR amplification of libraries was carried out using KAPA Hifi Polymerase. A post-PCR Agilent Bioanalyzer was used to quantify samples, followed by sample pooling and size-selection of pools using the LabChip XT Caliper. The multiplexed libraries were sequenced on the Illumina HiSeq 2000 for cohort 1 and HiSeq 4000 for cohorts 2-4 (75bp paired-ends). Sequenced data underwent initial analysis and quality control on reads as standard. The sequencing depth was similar across samples, with 90% of samples passing final QC (see below) having 87.2-129.2 million reads.

### Proteomics

Proteomics analysis was performed on cartilage samples from 103 patients (Supplementary Table 9). For

#### Cohort 1

All steps of protein digestion, 6-plex TMT labelling, peptide fractionation and LC-MS analysis on the Dionex Ultimate 3000 UHPLC system coupled with the high-resolution LTQ Orbitrap Velos mass spectrometer (Thermo Scientific), were previously described^9^. The sample preparation protocol formed the basis of processing for cohorts 2-4 using 10-plex TMT labelling and an Orbitrap Fusion Tribrid Mass Spectrometer (Thermo Scientific) with otherwise only minor alterations as described in the following.

#### Cohorts 2-4

##### Protein Digestion and TMT Labeling

The protein content of each sample was precipitated by the addition of 30 μL TCA 8 M at 4 °C for 30 min. The protein pellets were washed twice with ice cold acetone and finally re-suspended in 40 μL 0.1 M triethylammonium bicarbonate, 0.1% SDS with pulsed probe sonication. Equal aliquots containing at least 10 μg of total protein were reduced with 5 mM TCEP for 1h at 60 °C, alkylated with 10 mM Iodoacetamide and subjected to overnight trypsin (70 ng/μL) digestion. TMT 10-plex (Thermo Scientific) labelling was performed according to manufacturer’s instructions at equal amounts of tryptic digests. Samples were pooled and the mixture was dried with speedvac concentrator and stored at -20 °C until the peptide fractionation.

##### Peptide fractionation

Offline peptide fractionation was based on high pH Reverse Phase (RP) chromatography using the Waters, XBridge C18 column (2.1 x 150 mm, 3.5 μm) on a Dionex Ultimate 3000 HPLC system. Mobile phase A was 0.1% ammonium hydroxide and mobile phase B 100% acetonitrile, 0.1% ammonium hydroxide. The TMT labelled peptide mixture was dissolved in 100 μL mobile phase A, centrifuged and injected for fractionation. The gradient elution method at 0.2 mL/min included the following steps: 5 minutes isocratic at 5% B, for 35 min gradient to 35% B, gradient to 80% B in 5 min, isocratic for 5 minutes and re-equilibration to 5% B. Signal was recorded at 280 nm and fractions were collected every one minute. For cohort 4, peptide fractionation was performed on reversed-phase OASIS HLB cartridges at high pH and up to 9 fractions (10-25% acetonitrile elution steps) were collected for each set. The collected fractions were dried with SpeedVac concentrator and stored at -20 °C until the LC-MS analysis.

##### LC-MS Analysis

LC-MS analysis was performed on the Dionex Ultimate 3000 UHPLC system coupled with the Orbitrap Fusion Tribrid Mass Spectrometer (Thermo Scientific). Each peptide fraction was reconstituted in 40 μL 0.1% formic acid and a volume of 7 μL was loaded to the Acclaim PepMap 100, 100 μm × 2 cm C18, 5 μm, 100 Ȧ trapping column with the μlPickUp mode at 10 μL/min flow rate. The sample was then analysed with a gradient elution on the Acclaim PepMap RSLC (75 μm × 50 cm, 2 μm, 100 Å) C18 capillary column retrofitted to an electrospray emitter (New Objective, FS360-20-10-D-20) at 45 °C. Mobile phase A was 0.1% formic acid and mobile phase B was 80% acetonitrile, 0.1% formic acid. The gradient method at flow rate 300 nL/min was: for 90 min gradient to 38% B, for 5 min up to 95% B, for 13 min isocratic at 95% B, re-equilibration to 5% B in 2 min, for 10 min isocratic at 10% B. Precursors were selected with 120k mass resolution, AGC 3×10^5^ and IT 100 ms in the top speed mode within 3 sec and were targeted for CID fragmentation with quadrupole isolation width 1.2 Th. Collision energy was set at 35% with AGC 1×10^4^ and IT 35 ms. MS3 quantification spectra were acquired with further HCD fragmentation of the top 10 most abundant CID fragments isolated with Synchronous Precursor Selection (SPS) excluding neutral losses of maximum m/z 18. Iontrap isolation width was set at 0.7 Th for MS1 isolation, collision energy was applied at 55% and the AGC setting was at 6×10^4^ with 100 ms IT. The HCD MS3 spectra were acquired within 110-400 m/z with 60k resolution. Targeted precursors were dynamically excluded for further isolation and activation for 45 seconds with 7 ppm mass tolerance. Cohort 4 were analyzed at the MS2 level with a top15 HCD method (CE 40%, 50k resolution) and a maximum precursor intensity threshold of 5×10^7^ using the same MS1 parameters as above in a 360 min gradient.

### Genotyping

We used Illumina HumanCoreExome-12v1-1 for genoting cohort 1 and Illumina InfiniumCoreExome-24v1-1 for genotyping cohort 2-4 patients.

### Quantification of RNA levels

We used samtools v1.3.1^41^ and biobambam v0.0.191^42^ to convert cram to fasq files after exclusion of reads that failed QC. We applied FastQC v0.11.5 to check sample quality^43^ and excluded 9 samples (Supplementary Table 9).

We obtained transcript-level quantification using salmon 0.8.2^44^ (with --gcBias and --seqBias flags to account for potential biases) and the GRCh38 cDNA assembly release 87 downloaded from Ensembl [http://ftp.ensembl.org/pub/release-87/fasta/homo_sapiens/cdna/]. We used tximport^45^ to convert transcript-level to gene-level scaled transcripts per million (TPM) estimates, with estimates for 39037 genes based on Ensembl gene IDs.

We excluded 4 samples due to low mapping rate (<80%), 3 samples due to non-European ancestry recorded in the clinic, 18 samples due to low RIN (<5), 2 samples as duplicates, 8 samples due to abnormal gene read density plots (detected separately in cartilage and synovium for 3 cartilage and 5 synovium samples; all exclusions are listed in Supplementary Table 9).

The final gene expression dataset included 259 samples (Figure 1; 87 patients’ low-grade and 95 high-grade cartilage samples with 15,249 genes that showed counts per million (CPM) of ≥1 in ≥40 samples, and 77 patients’ synovium samples with 16,004 genes that showed CPM ≥1 in ≥20 samples).

### Quantification of protein levels

To carry out protein identification and quantification, we submitted the mass spectra to SequestHT search in Proteome Discoverer 2.1. The precursor mass tolerance was set at 30 ppm (Orbitrap Velos data, cohort 1) or 20 ppm (Fusion data, cohorts 2-4). For the CID spectra, we set the fragment ion mass tolerance to 0.5 Da; for the HCD spectra, to 0.02 Da. Spectra were searched for fully tryptic peptides with maximum 2 miss-cleavages and minimum length of 6 amino acids. We specified static modifications as TMT6plex at N-termimus, K and Carbamidomethyl at C; dynamic modifications included deamidation of N,Q and oxidation of M. For each peptide, we allowed for a maximum two different dynamic modifications with a maximum of two repetitions. We used the Percolator node to estimate peptide confidence. We set the peptide false discovery rate (FDR) at 1% and based validation on the q-value and decoy database search. We searched all spectra against a UniProt fasta file that contained 20,165 reviewed human entries. The Reporter Ion Quantifier node included a TMT-6plex (Velos data, cohort 1) or TMT-10plex (Fusion data, cohorts 2-4) custom Quantification Method with integration window tolerance at 20ppm or 15 ppm, respectively. As integration methods, we used the Most Confident Centroid at the MS2 or MS3 level. We only used peptides uniquely belonging to protein groups for quantification.

We excluded samples from 4 patients due to non-European ancestry (Supplementary Table 9). The final dataset included low-grade and high-grade cartilage samples each from 99 patients, with 4,801 proteins was observed in ≥30% of samples, and 1,677 proteins in all samples. To account for protein loading, abundance values were normalised by the sum of all protein abundances in a given sample, then log2-transformed and quantile normalised.

### Genotype analysis and quality control

Genotypes were called using GenCall (Illumina) and mapped to GRC37/hg19 using on-line tools (http://www.well.ox.ac.uk/~wrayner/strand/index.html). Quality control (QC) was carried out using the same method for both arrays. Briefly, we performed a pre-filtering step to exclude samples and variants with a call rate <90%. Sample QC included identity checks correlating the array genotypes to Fluidigm genotypes obtained at sample reception (no samples had a concordance <0.95). We excluded samples based on call rate <98%, heterozygosity distribution outliers performed using 2 different minor allele frequency (MAF) bins (≥1% MAF and <1% MAF) and sex discrepancies. We performed pairwise identity by descent (IBD) in PLINK^46, 47^ after filtering out variants with MAF <1% and carrying out linkage disequilibrium based pruning using R^2^ 0.2. We retained only patients with pairwise PI_HAT ≤0.2. To look at ethnicity we combined all patients from both arrays with data from the 1000 Genomes Project individuals^48^ . We included overlapping variants only and conducting IBD, as described above, followed by multidimensional scaling using PLINK. Visual ethnic outliers were excluded following examination of the first 2 components. Variants were excluded if call rate <98% and/or Hardy Weinberg p-value (pHWE) <1x10^-4^. The final datasets contained 12 patients and 534,694 variants and 99 patients and 527,717 variants for cohorts 1 and 2-4 respectively.

Prior to imputation, all genotypes were combined into a single dataset containing 111 patients and 504,235 overlapping variants. Further QC was performed to exclude any variants with strand, position and allele frequency differences compared to the HRC panel^49^ using a HRC preparation checking tool (http://www.well.ox.ac.uk/~wrayner/tools/; v4.2.7). The resulting dataset contained 111 patients and 389,511 variants. We imputed up to HRC panel (v1.1 2016) using the Michigan imputation server (https://imputationserver.sph.umich.edu/index.html)^50^ with Eagle2 (v2.3) phasing. Post-HRC imputation we used a post-imputation data checking program (http://www.well.ox.ac.uk/~wrayner/tools/Post-Imputation.html; v1.0.2) to visualise the results and we excluded variants with poor imputation quality (R^2^<0.3) and pHWE <1x10^-4^. We excluded two patients due to absence of RNA and protein data. The resulting final dataset contained 10,249,108 autosomal variants and 109 patients.

### Identification of *cis*-eQTLs and *cis*-pQTLs

We followed a similar method to GTEx^6, 7^.

#### cis-eQTLs

For each tissue, we included only genes with ≥1 count per million in at least 20% samples and we normalised between samples using TMM (weighted trimmed mean of M-values)^51^ implemented in edgeR^52^. To facilitate cartilage comparisons post-analysis, the previous 2 steps (exclusions of low expressed genes and the between sample normalisation) were performed with high-grade and low-grade cartilage samples combined. For each tissue separately, we then normalised across samples using an inverse normalisation transformation for each gene. To infer hidden factors associated with cohort, sequencing batch, or other technical differences, we applied Probabilistic Estimation of Expression Residuals (PEER)^53^ separately to each tissue (PEER C++ version with standard parameters from the R version, i.e. iteration=1000, bound=0.001, variance=0.00001, Alpha a = 0.001, Alpha b = 0.1, Eps a = 0.1, Eps b = 10). We used the GTEx modified version of FastQTL^54^ (https://github.com/francois-a/fastqtl; v6p) which allows for minor allele count filtering, reporting of minor allele frequency and calculation of FDR. We determined the transcription start site (TSS) for each gene using empirical transcript level expression information from synovium, high-grade and low-grade cartilage samples (see below) and defined the *cis*-mapping region to be 1Mb in either direction from the TSS. We restricted the analysis to variants with minor allele count of at least 10 in a given tissue. Nominal p-values for each gene-variant pair were based on linear regression, including 15 PEER factors for the given tissue, sex and genotype array as covariates. We then employed the adaptive permutation scheme with the --permute 1000 10000 option to generate empirical p-values. Genes with significant eQTLs (“eGenes”) were defined at the 5% Storey-Tibshirani False Discovery Rate (FDR) using the q-values generated from the empirical p-values^8^. For each eGene, significant eQTLs were defined as variants with nominal p-value below the nominal p-value threshold for that gene generated in FastQTL.

The normalised effect size (NES) of the eQTL is reported for the alternate allele according to GRC37/hg19.

#### cis-pQTLs

We followed a similar protocol as for *cis*-eQTL analysis. For low-grade and high-grade cartilage, we included 1677 proteins that were measured across all samples. We normalised across samples using an inverse normalisation transformation for each gene separately in each tissue. To account for possible technical variation, we used PEER^53^ (with parameters as for the eQTL analysis above) and included 26 PEER factors, sex and genotype array as covariates using the GTEx modified version of FastQTL (https://github.com/francois-a/fastqtl; v6p). We used the TSS established for the eQTL analysis, yielding a unique mapping for 1461 proteins, which were then taken forward. For each protein, we considered variants within a 1Mb region in either direction from the TSS, restricting further to minor allele count of 10 or higher. We then followed the same procedure as for *cis*-eQTLs to identify variant-protein pairs with significant *cis*-pQTL effects.

For both eQTLs and and pQTLs, we verified that the results were robust by carrying out a sensitivity analysis including patient age and osteoarthritis joint (knee or hip) as covariates in addition to the PEER factors, sex and array (Supplementary Note).

### Transcription start site (TSS) definition

To determine which transcript to use to define the TSS for each gene, we established the most abundant transcript for each gene. To have the same definition across all tissues, we analysed cartilage and synovium tissues jointly, considering 16 886 genes that passed quantification QC in at least one tissue (based on 15 249 genes in cartilage and 16 004 genes in synovium). For each transcript, we calculated the expression in each sample as scaled transcripts per million using tximport^45^. For each gene, we then obtained the most abundant transcript in each tissue in each patient, and calculated the proportion of samples in which each transcript was the most abundant. The transcript that was the most abundant in the largest proportion of samples was used to define the TSS for the gene. For genes in which more than one transcript was the most abundant, we chose one of the most abundant transcripts at random to define the TSS.

Across all genes, 47.9% had the same most abundant transcript in at least 90% of samples in both cartilage and synovium; 71% of genes had the same most abundant transcript in 60% of samples in cartilage and synovium. We mapped the most abundant transcript (using the ENST identifier) to GRCh37 using ftp://ftp.ensembl.org/pub/grch37/release-87/gtf/homo_sapiens/Homo_sapiens.GRCh37.87.chr.gtf.gz. Excluding 1940 transcripts with missing start or end positions (largely in patched genome build regions), and ∼500 transcripts mapped to chromosomes X, Y or mitochondrial DNA, we established the TSS for 13,180 autosomal genes included in the cartilage and 13,708 genes included in the synovium molQTL analysis.

### Differential gene regulation in low-grade and high-grade cartilage

To identify *cis*-eQTLs active exclusively in low-grade or high-grade cartilage, we used Meta-Tissue v0.5^55^ (downloaded from http://genetics.cs.ucla.edu/metatissue/index.html) which implements METASOFT^56^. The m-value calculated by METASOFT for gene-variant pair in each tissue provides a posterior probability (m value) of an effect in that tissue. Consequently, we aimed to identify eQTLs present in one tissue (defined as m>0.9), and absent in the other (defined as m<0.1). We note that there were no *cis*-eQTLs present in both tissues (m>0.9) with opposing direction of effect.

Meta-Tissue restricts covariates input to the same values for each patient across tissues, while different PEER covariates were provided for each tissue in the FastQTL analysis. Hence, for each tissue, we obtained residuals from regressing the normalised expression data on the 15 PEER factors, sex and array, then used the residuals as input for Meta-Tissue. We included genotype dosages based on both the low- and high-grade results for each analysis. We ran METASOFT using the default settings provided in the output script from Meta-Tissue. We only considered eQTLs that were identified in the FastQTL analysis in the appropriate tissue. To identify variants located in regulatory regions, we used Ensembl Variant Effect Predictor (http://grch37.ensembl.org/Homo_sapiens/Tools/VEP/).

### Colocalisation between molecular QTLs (molQTLs) and osteoarthritis GWAS associations

To examine colocalisation between molQTLs and GWAS associations, we used genome-wide summary statistics from the largest osteoarthritis meta-analysis to date, based on UK Biobank and arcOGEN data^5^. We analysed all 64 genome-wide significant signals using coloc^57^, separately for each tissue and omics level.

In the co-localisation analysis for each signal, we considered the region spanning 100kb either side of the index variants. If that region overlapped any genes with significant *cis*- eQTLs or *cis*-pQTLs, we extended the region to encompass all variants included in the molQTL analysis for these genes. To formally obtain a posterior probability for co-localisation, we used coloc.fast (https://github.com/tobyjohnson/gtx/blob/526120435bb3e29c39fc71604eee03a371ec3753/R/coloc.R), a Bayesian statistical test which implements the coloc^57^ method. We used the default settings for coloc.fast. We considered a 80% posterior probability of GWAS and molQTL shared association at a single variant (“PP4≥0.8”) to indicate evidence of colocalisation.

### Differential RNA expression between high-grade and low-grade cartilage

We tested differential expression of 15,249 genes between high-grade and low-grade cartilage using paired samples from 83 patients. To detect robust gene expression differences, we carried out analyses using different software packages as recommended in a landmark survey of best practices^58^, applying limma^59^, edgeR^60^, and DESeq2^61^. We also tested 5 analysis designs with different options to account for technical variation, including SVAseq^62^. In particular, we tested for differential expression using

(1) a paired analysis of intact and degraded samples (i.e. specifying patient ID as covariate),
(2) a paired analysis of intact and degraded samples, with 10 additional covariates accounting for technical variation identified by SVAseq^62^ ;
(3) a paired analysis of intact and degraded samples, with 10 RNA sequencing batches as covariates;
(4) an unpaired analysis of intact and degraded samples;
(5) an unpaired analysis of intact and degraded samples, with 19 additional covariates accounting for technical variation identified by SVAseq.

We tested for differential expression using the following R packages:

(i) limma^59^ (with lmFit and eBayes), after applying limma-voom^63^ to remove heteroscedasticity;
(ii) DESeq2^61^, separately with and without outlier filtering/replacement (minReplicatesForReplace=Inf, cooksCutoff=FALSE options);
(iii) edgeR^60^, using the likelihood ratio test (glmFit and glmLRT functions), and separately, using the F test (glmQLFit and glmQLFTest functions).
(iv) and elsewhere, we used Ensembl38p10 to identify genes with uniquely corresponding Ensembl gene ID and gene name (13,737 of 15,249 genes in the RNA data).
(v) each analysis design and method, we used a 5% False Discovery Rate (FDR) threshold to correct for multiple testing. This yielded 2,557 genes with significant differential expression between low-grade and high-grade cartilage across all analysis designs and testing methods (2,418 with uniquely corresponding Ensembl gene ID and gene name).

### Differential protein abundance between high-grade and low-grade cartilage

We performed differential analysis for 4,801 proteins that were measured in ≥30% of patients, applying limma^59^ to paired samples from 99 patients. Significance was defined at 5% FDR to correct for multiple testing, yielding 2,233 proteins with significant differential abundance (2,019 proteins with uniquely corresponding Ensembl gene ID and gene name). Paired samples from any patient were always assayed in the same multi-plex and we verified the results using a sensitivity analysis that explicitly accounts for technical variation to confirm that adjustment for patient effects was captured between-plex batch effects (Supplementary Note).

### Pathway associations for differences between high-grade and low-grade cartilage

To identify the biological processes with significant molecular differences between high-grade and low-grade cartilage, we carried out gene set enrichment analyses based on the differential expression (DE) on RNA, protein, and cross-omics levels. We tested for association of the differentially expressed (DE) genes on RNA and/or protein levels at 5% FDR, with robustness checks using more stringent FDR thresholds (1%, 0.5%, 0.1%). We restricted this analysis to genes with unique mapping between Ensembl gene ID and Gene Name in Ensembl38p10, and Ensembl gene ID and Entrez ID in HUGO (https://www.genenames.org/ accessed 05/03/2018; 13,094 genes measured on RNA level, 4,390 genes measured on protein level, 4,387 genes measured on both RNA and protein levels).

We applied Signalling Pathway Impact Analysis (SPIA)^64^ to test for association with KEGG signaling pathways. SPIA combines enrichment p-values with perturbation impact on the pathway based on log-fold differences of the DE genes; perturbation p-values are obtained by bootstrapping. Enrichment and perturbation p-values were combined using a normal inversion method which only gives low p-values when both over-representation and pathway impact p-values are low (function option combine=“norminv”). Significance of pathway association was defined as a threshold of 5% FDR applied to the combined p-values in each analysis. For the analysis of genes DE on both RNA and protein levels, we carried out tests using the log-fold differences from the RNA data (based on the limma analysis with paired samples and SVAseq covariates), and separately, from the protein data.

We also tested enrichment in Gene Ontology terms using GOseq^65^, separately for genes with higher or lower expression in high-grade compared to low-grade cartilage. We accounted for gene length (pwf function options “hg19” and “geneSymbol”). Significance was defined as a threshold of 5% FDR in each analysis. The results showed broad agreement with the results of the SPIA analysis (Supplementary Table 3).

### Identification of genes with osteoarthritis GWAS gene-level association

From the recent UK Biobank and arcOGEN GWAS meta-analysis^5^, we obtained the results of a gene-level analysis for each of the four osteoarthritis phenotypes (self-reported plus hospital diagnosed, hospital diagnosed knee or hip, hospital diagnosed knee, hospital diagnosed hip), as described in the GWAS paper. Briefly, this analysis used MAGMA v1.06^66^ and was based on the mean SNP log-p-value in the gene, accounting for LD.

To calculate the effective number of tests across phenotypes, we calculated the correlation matrix between the gene p-values for the four osteoarthritis phenotypes, and obtained the eigenvalues of this matrix. The effective number of tests *N_eff_* for phenotypes was then calculated as

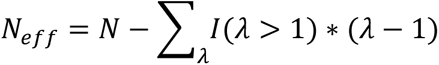

where N=4 is the number of phenotypes, and *λ* denotes the eigenvalues. Across the Pearson and Spearman correlation matrices, we obtained *N_eff_*<2.65. With 18449 genes per phenotype, the significance threshold for gene-level p-values across genes and phenotypes was thus set as 0.05/(18449*2.65)=1.02x10^-6^.

After accounting for the effective number of tests across phenotypes and genes using a Bonferroni correction, 320 of 18,449 genes showed significant association with at least one phenotype. Of these genes, 238 genes were compared between low-grade and high-grade cartilage on at least one omics level and had uniquely corresponding Ensembl gene ID and gene name.

### Sample clustering

For RNA data, we normalised each tissue separately, using limma-voom^63^ to remove heteroscedasticity from scaled TPM values. Especially when considering clustering within a particular tissue, technical effects can influence the results. Consequently, for RNA data, we applied pSVA^67^ (designed specifically to preserve biological heterogeneity for clustering) to the voom-normalised data, separately within low-grade cartilage, high-grade cartilage, and synovium, using the RNA sequencing batches to remove technical variation (post-QC, 9 batches for low-grade cartilage, 10 batches for high-grade cartilage, 8 batches for synovium). After this step, we obtained residuals from linear regression of the post-pSVA-data on an artificial variable combining RNA sequencing batches with clinical batches (i.e. batches in which samples were sent to the sequencing facility, with paired samples always sent in the same batch; in total, 12 sequencing and clinical batches for low-grade cartilage, 13 for high-grade cartilage, 11 for synovium).

For the proteomics data, we regressed out batches from the log2-transformed normalised abundance values. We then quantile normalised the residuals as analogous step to the differential expression analysis; all analyses were also done without quantile normalisation, with no appreciable difference in results (we did not detect any clustering within proteomics tissue, see Supplementary Figure 3d-e).

For each tissue and omics level, we applied ConsensusClusterPlus^68^, a consensus clustering method that splits samples into a discrete number of groups, so that samples within a group are more similar to each other than to samples outside the group. We used standard settings, with scaled gene expression or protein abundance values, a Euclidean distance between samples, and a hierarchical clustering algorithm (options innerLinkage=“average”, finalLinkage=“average”, corUse=“complete.obs”, clusterAlg=“hc”). The maximum number of clusters was 10, with 1000-fold re-sampling for 80% of samples and a fixed seed for reproducibility (options maxK=10, reps=1000, pItem=0.8, pFeature=1, seed=1262118388.71279). The final number of clusters was chosen based on the Consensus Cumulative Distribution Function plots, the Delta Area Plot, and a visual investigation of the Consensus Matrices, as advised in the manual. Results were confirmed via additional analysis using a distance metric based on Pearson correlation.

We checked that clustering was not associated with patient cohort (chi-square test, p>0.99 in synovium and low-grade cartilage) nor with batches samples were sent and sequenced (chi-square test, *P*>0.96 in synovium and low-grade cartilage).

### Differential gene expression between tissue clusters

To follow up the clustering results for low-grade cartilage and synovium, we tested gene differential expression between sets of samples based on cluster assignment (applying limma to the normalised expression values underlying the clustering, i.e. gene expression after voom, pSVA, and regression of batch covariates). The differential expression analysis was followed up by gene set enrichment analyses using SPIA and GOseq, with 8 gene differential expression FDR thresholds to assess robustness of the association (5%, 0.5%, 5x10^-3^, …, 5x10^-7^). In each analysis, gene set association was defined at the 5% FDR threshold. As before for GOseq, genes with positive and negative log-fold-difference between clusters were analysed separately.

### Differential gene expression between high-grade and low-grade cartilage by low-grade cartilage clusters

To test whether differences between high-grade and low-grade cartilage depended on the low-grade cartilage sample cluster, we carried out differential expression analyses separately for patients in cartilage-Cluster1, and in cartilage-Cluster2. We used limma-voom and a paired-sample design as in the main differential expression analysis. We then computed the Spearman correlation of gene log-fold differences in the main and cluster-specific analyses using cor.test in R. We repeated the correlation analysis for the genes with significant differences in the all-patient analysis.

### Multi-omics factor analysis (MOFA) and correspondence to sample clustering

To test for patient heterogeneity using a method that can detect both discrete clustering and a continuous spectrum of variation, we used multi-omics factor analysis (MOFA)^14^. MOFA can integrate data across omics levels and across tissues to discover hidden factors that represent drivers of variability between samples or patients. MOFA was run i) jointly on all RNA and protein data; ii) jointly on RNA data across all three tissues; iii) on RNA and protein data within each tissue. MOFA identifies a factor score for each sample or patient, calculates the variance explained by each factor in each omics level and tissue, calculates weights of genes on each factor from each omics level and tissue, carries out a gene set enrichment for each factor in each omics level and tissue based on gene weights. All analyses were restricted to genes and proteins with unique correspondence between Ensembl gene ID and gene name as above. The technical parameters applied were gaussian likelihoods, 5000 iterations, a maximum of 100 factors, and dropping factors that explain less than 5% during training, with fixed seed 20180613 for reproducibility.

We further investigated the correspondence between the discrete clusters identified by ConsensusClusterPlus and the spectrum of variation identified by the MOFA as follows. We tested the gene expression differences between subsets of the discrete low-grade tissue clusters at a MOFA low-grade factor 1 threshold of 0, which corresponded most closely to the cluster assignment, with consistent assignment for 84% of patients in the first cluster (38 of 45 with score >0) and 83% of patients in the second cluster (35 out of 42 with score <0). We analysed gene expression differences between the 38 and 7 samples in cartilage-Cluster1 with MOFA factor 1 values above and below 0, respectively. Analogously, we analysed gene expression differences between the 7 and 35 samples in cartilage-Cluster2 with MOFA low-grade factor 1 values above and below 0, respectively. We applied limma to the normalised gene expression values underlying the clustering, i.e. gene expression after voom, pSVA, and regression of batch covariates.

Second, we calculated the Spearman correlation between gene weights from the RNA data on the MOFA low-grade factor 1 and the log-fold differences between the two low-grade cartilage clusters using the cor.test function in R.

For synovium, we carried out a Spearman correlation analysis between the gene weights on the MOFA synovium factors 1 and 2, and the log-fold differences between synovium-Cluster1 and synovium-Cluster2, as well as between the synovium subclusters synovium-Cluster1a and synovium-Cluster1b, and separately, synovium-Cluster2a and synovium-Cluster2b.

### Construction of classifier to reflect sample clustering

We applied PAMR^15^, a soft-thresholding centroid-based method, to identify a smaller subset of genes which could distinguish the low-grade cartilage clusters. We first computed silhouette scores for each low-grade sample based on the clustering, to calculate how similar each sample is to samples in the same cluster versus the other cluster For each sample *i* in cartilage-Cluster1, we obtained the average dis-similarity to other samples in cartilage-Cluster1 (written *j* ∈ *C*_1_, *j* ≠ *i*) as 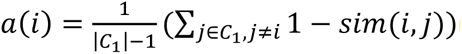 where *sim*(*i, j*) is the similarity between samples *i* and *j* in cartilage-Cluster1 computed by ConsensusCluster, and |*C*_1_| is the number of samples in cartilage-Cluster1. We then obtained the average dis-similarity of sample *i* to samples in cartilage-Cluster2 as 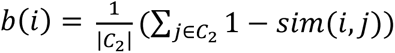 The silhouette score for sample *i* was then calculated as 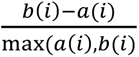 We proceeded analogously for samples in cartilage-Cluster2.

To train a classifier, we considered all samples with silhouette score >0.2 (removing 1 sample in low-grade cartilage-Cluster1, 3 samples in low-grade cartilage-Cluster2). We also restricted the analysis to 1,063 genes with high expression levels, obtained as median scaledTPM value of at least 5,000 across all low-grade cartilage samples. We applied pamr to the 1063 genes and 83 samples, setting the seed to 20180927 for all random components. We applied the function pamr.adaptthresh (with options ntries = 100, reduction.factor = 0.9) to identify thresholding scales for the classifier training. We then used the pamr.train function to train a classifier, and the pamr.cv function to examine the classifier error rates in cross-fit validation within the training data. We identified the minimum error rate to be reached with a threshold of 6.666334, yielding 7 genes and a cross-validation error rate of 0.07. We then used the pamr.predict function to predict cartilage-Cluster1 and cartilage-Cluster2 probabilities for all 87 low-grade samples, with an agnostic prior setting of 0.5 for both clusters (options type=“posterior”,prior=c(0.5,0.5)). We also calculated Spearman correlations for the PAMR cluster probabilities and MOFA Factor1 for low-grade cartilage.

### Validation of clustering classifier

We obtained RNA expression data from low-grade cartilage tissue of 60 knee osteoarthritis patients undergoing joint replacement (27 women, 33 men, age range 63-85 years), sequenced on Illumina HiSeq 2500, with transcript quantification using kallisto and quality control as described previously^13^. We obtained the gene-level expression data from Github (file txi.RData on github.com/soulj/OAStratification, accessed 03/10/2018), with sample batch information provided in a separate file (patientDetails_all_withMed.csv). We then carried out further steps to harmonise data processing with our approach. First, we used tximport to transform the expression data to scaled transcript per million (scaledTPM) levels. We then applied the voom function in the limma R package to remove heteroscedasticity, followed by applying pSVA to remove batch effects (based on the known batches as listed in patient details). As for the data used in this study, we calculated residuals from a linear regression of post-pSVA data on batches, and used these expression residuals as data post batch effect removal.

We applied the pamr.predict function to predict cartilage-Cluster1 and cartilage-Cluster2 probabilities for all 60 samples using the trained 7-gene classifier, with an agnostic prior setting of 0.5 for both clusters. We also applied MOFA (with the same parameters and options as above) to the data post batch effect removal. Finally, we calculated Spearman correlations for the PAMR cluster probabilities and MOFA factor 1.

The original publication also included a division of samples into 2 groups using non-negative matrix factorisation based on known biological networks. The group assignment was also provided on Github (file NetNMF_R2_L25.RData). This assignment was compared to a cluster assignment based on PAMR 7-gene classifier posterior probabilities.

### Associations between tissue cluster assignment and clinical data

We tested for association between low-grade cartilage dichotomous cluster assignment (high-inflammation cartilage-Cluster1 versus low-inflammation cartilage-Cluster2) and clinical characteristics using a generalised linear model (via the glm function in R with option family=”binomial”). To consider the association of tissue clusters with drug prescription, drugs were grouped by pharmacological mechanism into 58 categories by two clinical experts (AF & JMW). We restricted the analysis to 9 drug categories, each with at least 20 patients who were also assigned a low-grade cartilage or synovium cluster (Supplementary Table 7).

To identify the number of effective tests across clinical characteristics (age, height, weight, BMI, ASA grade, number of drugs taken) and the 9 drug categories, we calculated pairwise correlations across patients using pairwise complete observations. We then obtained and eigenvalues of the correlation matrix, and calculated the effective number of tests *N_eff_* for clinical characteristics as

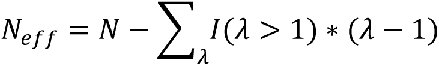

where N=16 is the number of characteristics tested, and *λ* denotes the eigenvalues. Across the Pearson and Spearman correlation matrices, we obtained *N_eff_*<10.54. We thus used a Bonferroni-corrected threshold of *P*<0.05/10.54=0.0047 to define significance.

As a sensitivity analysis, we tested for association of low-grade cartilage dichotomous cluster assignment (high-inflammation cartilage-Cluster1 versus low-inflammation cartilage-Cluster2) using a generalised linear model (via the glm function in R with option family=”binomial”), accounting for sex or sex and age (Supplementary Note).

For association with synovium cluster assignment, we carried out analogous tests for the two clusters, with the same Bonferroni-corrected significance threshold.

## ConnectivityMap analysis

To identify opportunities for drug repurposing, we used ConnectivityMap^27^ to identify compounds and perturbagen classes (PCL) that could possibly reverse the differences identified between high-grade and low-grade cartilage. Using the online interface clue.io (accessed 03/03/2019), we submitted the 148 genes with significantly higher expression on both RNA and protein level to calculate a “tau” connectivity score to gene expression signatures experimentally induced by various perturbations in 9 cell lines. A positive tau score indicates similarity between the gene expression signature of a perturbation and the submitted query (i.e. up-regulation of the genes with higher expression in high-grade compared to low-grade cartilage). A negative tau score indicates that gene expression signature of a perturbation opposes the submitted query (i.e. down-regulation of the genes with higher expression in high-grade compared to low-grade cartilage). Recommended thresholds for further consideration of results are tau of at least 90, or below -90, respectively (https://clue.io/connectopedia/connectivity_scores, accessed 03/03/2019). A total of 2837 compound and 171 PCL perturbations were evaluated in clue.io. We shortlisted perturbations where both the summary tau and the median tau across cell lines were higher than 90 or lower than -90 for perturbagen classes, with more conservative thresholds of higher than 95 or lower than -95 for compounds. The clue.io platform also contained perturbation data from 3799 gene knock-down and 2160 over-expression experiments (with 2111 genes in both, i.e. 3848 genes total). These data were used to shortlist genes where both the summary and median tau were higher than 95 or lower than -95.

## Generation and phenotyping of mouse mutant lines

Targeted genetically-modified lines with deletion alleles for mouse orthologues of 7 shortlisted human genes were generated by the Wellcome Sanger Institute as part of the International Knockout Mouse Consortium (IMPC), on the same C57BL/6N:C57Bl/6NTac background (Supplementary Figure 8a). Targeted constructs for *Clic3*, *Crip1*, *Htra3*, *Matn4* and *Pdlim1* and *Sqrdl* were produced by CRISPR endonuclease deletion. The *Cpt1a* mutant allele (tm1.2) was produced by replacing the critical exon with a LacZ/neomycin reporter cassette, which was subsequently removed by Flp recominbase via FRT sites flanking the construct^69^. We confirmed human-mouse gene orthology as high-confidence one-to-one orthologues for 5 genes (*CLIC3*, *CPT1A*, *HTRA3*, *MATN4*, *PDLIM1*) and high-confidence many-to-one orthologues for the other two human genes (*CRIP1*, *SQOR*) in Ensembl38p10. All mutant and wild type (WT) mice were 16-week old male mice. For each mutant line, three mice were phenotyped (with four mice for Htra3) and compared to a 100-sample WT reference range for each parameter. While the number of mice was limited, previous extensive validation of the joint phenotyping pipeline showed that measurements were robust and strong phenotypic outliers could be detected^37^.

## Joint phenotype analysis for the mouse mutant lines

The joint phenotype the left hindlimb for each line was analysed using a rapid-throughput pipeline, as described in detail elsewhere^37^. Tibial articular cartilage volume (mm^3^), and median and maximum cartilage thickness (mm) were evaluated using micro-computed tomography with the aid of a contrast medium to segment out soft tissue (Iodine contrast-enhanced microCT, ICE-µCT). Subchondral bone mineral density (mg/hydroxyapatite/mm^3^), bone volume per tissue volume (BV/TV, %), trabecular number and trabecular thickness (mm) were evaluated in the underlying subchondral region. Aerial bone mineral content (median grey level) was quantified using subchondral X-ray microradiography (scXRM). Articular cartilage surface damage area (%) on the tibial plateaux was quantified using scanning electron microscopy of resin replicas of the articulating surfaces (Joint Surface Replication, JSR). The medial (MTP) and lateral (LTP) plateaux were assessed separately. The efficacy of these novel joint phenotyping methods in detecting signs of osteoarthritis in the mouse knee was confirmed by phenotyping limbs from mice in which post-traumatic osteoarthritis was provoked by DMM (Destabilisation of the Medial Meniscus) surgery^70^. These methods detected significant signs of osteoarthritis in the medial compartment of DMM-operated knees compared to sham, correlated to a high OARSI score^37^.

As the 18 phenotypes are not independent, we calculated the number of effective tests within each line using the information from the background mice as follows. We calculated the correlation matrix between the 18 phenotypes based on the 100 background mice, and obtained the eigenvalues of this matrix. The number of effective tests *N_eff_* was then calculated as

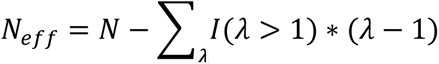

where *N*=18 is the number of parameters, and *λ* denotes the eigenvalues. Across the Pearson and Spearman correlation matrices, we obtained *N_eff_*<8.8. Therefore, applying a Bonferroni correction for the effective number of tests, line-wise significance for the differences between the background mice and each of the mouse mutant lines was defined as *P*<0.00568.

## Data Availability

The RNA sequencing data reported in this paper have been deposited to the EGA (accession numbers EGAS00001002255 (https://www.dev.ebi.ac.uk/ega/studies/EGAS00001002255), EGAD00001003355, EGAD00001003354, EGAD00001001331). The proteomics data reported in this paper have been deposited to PRIDE (accession numbers PXD014666, PXD006673, PXD002014). The genotype data reported in this paper have been deposited to the EGA (accession numbers EGAD00010001746, EGAD00010001285, EGAD00010001292, EGAD00010000722).

## Code Availability

All software used in this study is available from free repositories or from manufacturers as referenced in the Methods section.

## Supplementary Information

### Supplementary Note

This file contains additional notes on the molecular QTL analysis, the sensitivity analysis for differential protein abundance between high-grade and low-grade cartilage, examples of genes with cross-omics differences between high-grade and low-grade cartilage, details of mouse models and the identification of abnormal joint phenotypes, pathway associations for differences between low-grade and high-grade cartilage, and patient stratification.

### Supplementary Figures

**Supplementary Figure 1. Molecular QTLs in osteoarthritis disease tissue**

(a) eQTL overlap between tissues, for a total of 1,891 genes with a least one eQTL (left) and 219,709 eQTL gene-variant pairs (right). 49% of detected eQTLs are not tissue-specific.
(b) High correlation of eQTL normalized effect sizes (NES) between low-grade and high-grade cartilage. Inset: Spearman correlation ρ=0.94 between NES effect sizes across all eQTLs.
(c) High correlation of eQTL normalized effect sizes (NES) between low-grade cartilage and synovium (left), and between high-grade cartilage and synovium (right). Inset: Spearman correlation between NES effect sizes across all eQTLs.
(d) pQTL overlap between tissues, for a total of 38 genes with a least one pQTL (left) and 3,211 pQTL protein-variant pairs (right).
(e) High correlation of pQTL NES between low-grade and high-grade cartilage. Inset: Spearman correlation between NES effect sizes across all eQTLs.
(f) Plots of GWAS and low-grade cartilage eQTL p-values for regions surrounding rs10502437 and rs12901372. For both GWAS signals, we observed colocalisation with eQTLs in both low-grade and high-grade cartilage, and the plots for high-grade cartilage are shown in **Figure 2d**. Each plot shows 1Mb region centered around the GWAS index SNP (purple); each point represents a genetic variant. Top panels show GWAS p-values, bottom panels QTL p-values for the indicated gene. Here and in **Figure 2d**, LD between variants was calculated using UK Biobank. PP4: posterior probability for colocalisation.

**Supplementary Figure 2. Molecular differences between low-grade and high-grade cartilage**

(a) RNA- and protein-level log-fold differences for all genes measured on both omics levels. The x-axis shows gene expression differences between low-grade and high-grade cartilage as quantified by limma in an analysis including the technical covariates as identified by pSVA as well as pairing samples from the same patients (see **Methods**). The genes highlighted black or red were significant on both RNA- and protein-level, see also **Figure 3b**. The correlation of effect sizes across all genes was also significant (Pearson *r*=0.32, *P*<10^-10^).
(b) Signalling Pathway Impact Analysis (SPIA) identified biological pathways associated with high-grade/low-grade differences. All pathways shown are activated in high-grade compared to low-grade cartilage. Enrichment p: p-value from over-representation analysis of genes; Perturbation p: p-value for perturbation of the pathway based on gene log-fold-differences; Combined p: p-value from combining enrichment and perturbation p-values. Pathways with significant results at 5% FDR based on RNA-level changes are shown.

**Supplementary Figure 3. Cluster consensus and tracking plots for the clustering analysis of samples within tissues**

(a) Cluster consensus plots for clustering in low-grade cartilage, synovium, and high-grade cartilage based on RNA data. The x-axis shows the number k of clusters, the y-axis the cluster consensus value (higher values showing stronger clustering). For clustering in low-grade cartilage and synovium, but not high-grade cartilage, the cluster consensus value is above 0.8 for both clusters when k=2.
(b) High-grade cartilage tissue samples from patients do not show a separation into two clusters by ConsensusCluster analysis based on RNA data (cluster consensus value <0.8 for at least one cluster).
(c) Cluster tracking plots for low-grade cartilage and synovium samples based on RNA data. Each column is a sample, coloured by the cluster assignment when separating samples into k=2,…,10 clusters (k values in rows).
(d) Low-grade cartilage tissue samples from patients do not show a separation into clusters by ConsensusCluster analysis based on protein data. k: number of clusters.
(e) High-grade cartilage tissue samples from patients do not show a separation into clusters by ConsensusCluster analysis based on protein data. k: number of clusters.

**Supplementary Figure 4. Gene expression analysis of the clusters identified in low-grade cartilage and synovium tissue and correspondence between synovium clustering and MOFA results**

(a) Gene expression differences between low-grade and high-grade cartilage do not depend on the low-grade cartilage cluster. Plots show log-fold differences for all genes based on the analysis of all patients (x-axis) versus log-fold differences for all genes based on the analysis of all patients with low-grade cartilage in only one of the two clusters (y-axis). In each within-cluster analysis, 99% of the genes significant in the all-patient analysis had the same direction of effect. Inset: Spearman correlation of log-fold-differences, *P*<1.0x10^-10^.
(b) Gene expression differences between the synovium sub-clusters within each cluster are highly correlated. Plot shows log-fold differences of each gene in the comparison of sub-clusters within the larger (x-axis) and smaller (y-axis) cluster. Over 99% of the genes with significant differences between synovium-Cluster1a and synovium-Cluster1b also had directionally concordant differences between synovium-Cluster2b and synovium-Cluster2a, and over 80% were also significant at 5% FDR, and vice versa (i.e. genes with higher expression in synovium-Cluster1a compared to synovium-Cluster1b also had higher expression in synovium-Cluster2b compared to synovium-Cluster2a).
(c) An analysis of synovium samples using MOFA identifies a continuous spectrum of variation between samples. This variation corresponds well to the clustering: MOFA synovium factor 1 captures differences between sub-clusters, while synovium factor 2 captures variation between clusters.

**Supplementary Figure 5. Multi-omics factor analysis (MOFA) RNA gene weights are correlated with gene expression differences between tissue clusters**

(a) Correlation between MOFA low-grade cartilage Factor 1 gene weights for RNA data and gene expression differences between low-grade cartilage clusters. Inset: Spearman correlation, *P*<10^-15^. Genes with significant differential expression between low-grade and high-grade cartilage are coloured red.
(b) Gene expression differences between low-grade cartilage samples in the same clusters, divided by MOFA low-grade cartilage factor 1 values >0 and <0, are correlated with gene expression differences between clusters. Inset: Spearman correlation, *P*<10^-10^. Left shows results for low-grade cartilage-Cluster1, right for cartilage-Cluster2. Genes with significant differential expression between low-grade and high-grade cartilage are coloured red.
(c) Correlation between MOFA synovium Factor 1 and 2 gene weights for RNA data and gene expression differences between synovium clusters and subclusters. Inset: Spearman correlation, *P*<10^-15^.

C-Cluster: cartilage-Cluster.

**Supplementary Figure 6. PAMR 7-gene low-grade cartilage classifier performance**

PAMR diagnostic plots for a classifier of low-grade cartilage based on RNA. Left: Sample classification error based on the PAMR internal threshold and the corresponding number of genes in the classifier. The top panel shows the overall error estimate, the bottom panel error rates separately for cartilage-Cluster1 and cartilage-Cluster2. The optimal selection as used in the paper included 7 genes and an internal threshold of 6.67 (vertical line). Right: False Discover Rate (FDR) for between-cluster differences for the genes in the classifier as calculated by PAMR. C-Cluster: low-grade cartilage-Cluster.

**Supplementary Figure 7. Mouse models of implicated genes display osteoarthritis-relevant abnormal joint phenotypes.**

(a) Overview of the abnormal mouse joint phenotypes displayed by mouse models for 7 genes with differential expression between low-grade and high-grade cartilage. For each of the genes (in rows), 9 phenotypes were assayed on the lateral tibial plateau (LTP) and medial tibial plateau (MTP). For each joint parameter (in columns), the plot shows the ratio of the mean value of each mutant strain to the mean value of the wild-type background strain, where the differences were significant (*P*<0.05). Borders around boxes show phenotype difference significant after multiple-testing correction for the effective number of tests within each line (*P*<0.00568). BMC: bone mineral content; BMD: bone mineral density; vol: volume. Technical details of each mouse mutant line and plots of all individual values for abnormal phenotypes at *P*<0.05 see Supplementary Figure 8.
(b) Iodine contrast enhanced micro computed tomography (ICE-µCT) detects differences in articular cartilage volume and thickness (red volumes), and subchondral bone morphology (blue volumes); scale bar 100µm. *Matn4^-/-^* mice show decreased subchondral BV/TV (bone volume/tissue volume) and trabecular thickness (black arrow) compared to wild-type controls, whereas *Pdlim1^-/-^* mice display increased articular cartilage thickness (black arrow).
(c) Joint surface replication (JSR) detects damage to the articulating surfaces of the tibial plateaux; scale bar 100um. *Htra3^-/-^* mice show increased articular cartilage surface damage (black arrows, C) compared to wild-type controls.
(d) Subchondral X-ray microradiography (scXRM) detects changes in BMC within the subchondral region. White boxes represent the subchondral regions analysed; scale bar 1mm. *Matn4^-/-^* mice show decreased subchondral BMC compared to wild-type controls.
(e) All abnormal phenotypes displayed by *Matn4^-/-^*, *Pdlim1^-/-^* and *Htra3^-/-^* (*P*<0.00568). The boxplots show phenotype values for 100 mice from the background strain, with error bars for the 25%-75% interquartile range. Red diamonds: phenotypes of mutant mice. BV/TV: bone volume per tissue volume.

**Supplementary Figure 8. Details of mouse mutant lines and additional abnormal joint phenotypes displayed by mouse models of implicated genes**

(a) Technical details for each mouse mutant line, including targeting method and allele name.
(b) The figure shows all abnormal phenotypes displayed at *P*<0.05 which were not shown in the main figure. The boxplots show phenotype values for 100 mice from the background strain, with error bars for the 25%-75% interquartile range. Red diamonds: phenotypes of the mouse lines.

### Supplementary Tables

**Supplementary Table 1. Detailed list of differential eQTLs, i.e. variants with high posterior probability for presence of eQTL effect in high-grade cartilage (m>0.9) and low for presence in low-grade cartilage (m<0.1), or vice versa.**

**Supplementary Table 2. Genes with significant cross-omics differences between high-grade and low-grade cartilage.**

**Supplementary Table 3. Pathways and gene sets associated with significant RNA-level and/or protein-level differences between high-grade and low-grade cartilage.**

**Supplementary Table 4. Genes with significantly different expression profiles between high-and low-grade cartilage that were also found to be associated with genetic risk of osteoarthritis in a recent GWAS (multiple-testing corrected significance threshold of P<1.02x10^-6^).**

**Supplementary Table 5. Pathways associated with gene expression differences between low-grade cartilage clusters or between synovium clusters.**

**Supplementary Table 6. Expression differences between low-grade cartilage clusters for genes highlighted in previous cartilage clustering analyses.**

**Supplementary Table 7. Full association results between patient clinical characteristics and low-grade sample cluster assignment or synovium cluster assignment, including individual drugs assigned to drug classes.**

The nine drug classes which were tested for association are shown (see **Methods).**

**Supplementary Table 8. Comparison of perturbations by compounds, gene knockdown or overexpression, to differences between high-grade and low-grade cartilage.**

The comparison was based on data from ConnectivityMap and genes with higher expression in high-grade than in low-grade cartilage on both RNA- and protein-level, see **Methods.**

**Supplementary Table 9. List of all assayed patient tissue samples with detailed information including cohort, batch, and quality control exclusions.**

## References

1. Vos, T. et al. Global, regional, and national incidence, prevalence, and years lived with disability for 310 diseases and injuries, 1990–2015: a systematic analysis for the Global Burden of Disease Study 2015. Lancet 388, 1545–1602, doi:10.1016/S0140-6736(16)31678-6 (2016).

2. Mobasheri, A. et al. The role of metabolism in the pathogenesis of osteoarthritis. Nat. Rev. Rheumatol. 13, 302, doi:10.1038/nrrheum.2017.50 (2017).

3. Murphy, L. et al. Lifetime risk of symptomatic knee osteoarthritis. Arthritis Care Res. 59, 1207–1213, doi:10.1002/art.24021 (2008).

4. Murphy, L. B. et al. One in four people may develop symptomatic hip osteoarthritis in his or her lifetime. Osteoarthr. Cartil. 18, 1372–1379, doi:10.1016/j.joca.2010.08.005 (2010).

5. Tachmazidou, I. et al. Identification of new therapeutic targets for osteoarthritis through genome-wide analyses of UK Biobank data. Nat. Genet. 51, 230–236, doi:10.1038/s41588-018-0327-1 (2019).

6. Brown, A. A. et al. Predicting causal variants affecting expression by using whole-genome sequencing and RNA-seq from multiple human tissues. Nat. Genet. 49, 1747–1751, doi:10.1038/ng.3979 (2017).

7. GTEx Consortium et al. Genetic effects on gene expression across human tissues. Nature 550, 204–213, doi:10.1038/nature24277 (2017).

8. Storey, J. D. & Tibshirani, R. Statistical significance for genomewide studies. Proc. Natl. Acad. Sci. U.S.A. 100, 9440–9445, doi:10.1073/pnas.1530509100 (2003).

9. Steinberg, J. et al. Integrative epigenomics, transcriptomics and proteomics of patient chondrocytes reveal genes and pathways involved in osteoarthritis. Sci. Rep. 7, 8935, doi:10.1038/s41598-017-09335-6 (2017).

10. Karlsson, C. et al. Genome-wide expression profiling reveals new candidate genes associated with osteoarthritis. Osteoarthr. Cartil. 18, 581–592, doi:10.1016/j.joca.2009.12.002 (2010).

11. Ramos, Y. F. M. et al. Genes involved in the osteoarthritis process identified through genome wide expression analysis in articular cartilage; the RAAK Study. PLOS ONE 9, e103056, doi:10.1371/journal.pone.0103056 (2014).

12. Fernández-Tajes, J. et al. Genome-wide DNA methylation analysis of articular chondrocytes reveals a cluster of osteoarthritic patients. Ann. Rheum. Dis. 73, 668–677, doi:10.1136/annrheumdis-2012-202783 (2014).

13. Soul, J. et al. Stratification of knee osteoarthritis: two major patient subgroups identified by genome-wide expression analysis of articular cartilage. Ann. Rheum. Dis. 77, 423–423, doi:10.1136/annrheumdis-2017-212603 (2018).

14. Argelaguet, R. et al. Multi-Omics Factor Analysis—a framework for unsupervised integration of multi-omics data sets. Mol. Syst. Biol. 14, e8124, doi:10.15252/msb.20178124 (2018).

15. Tibshirani, R., Hastie, T., Narasimhan, B. & Chu, G. Diagnosis of multiple cancer types by shrunken centroids of gene expression. Proc. Natl. Acad. Sci. U.S.A. 99, 6567–6572, doi:10.1073/pnas.082099299 (2002).

16. Murphy, G. & Lee, M. H. What are the roles of metalloproteinases in cartilage and bone damage? Ann. Rheum. Dis. 64, iv44–iv47, doi:10.1136/ard.2005.042465 (2005).

17. Tanaka, T., Narazaki, M. & Kishimoto, T. IL-6 in inflammation, immunity, and disease. Cold Spring Harb. Perspect. Biol., doi:10.1101/cshperspect.a016295 (2014).

18. López-Boado, Y. S., Tolivia, J. & López-Otín, C. Apolipoprotein D gene induction by retinoic acid is concomitant with growth arrest and cell differentiation in human breast cancer cells. J. Biol. Chem. 269, 26871–26878, &lt;http://www.jbc.org/content/269/43/26871.abstract> (1994).

19. Shepherd, C. et al. Functional characterization of the osteoarthritis genetic risk residing at *ALDH1A2* identifies rs12915901 as a key target variant. Arthritis Rheumatol. 70, 1577–1587, doi:10.1002/art.40545 (2018).

20. Styrkarsdottir, U. et al. Severe osteoarthritis of the hand associates with common variants within the *ALDH1A2* gene and with rare variants at 1p31. Nat. Genet. 46, 498–502, doi:10.1038/ng.2957s (2014).

21. Owens, William D., M.D., Felts, James A., M.D. & Spitznagel, Edward L., Ph.D. ASA physical status classifications: A study of consistency of ratings. Anesthesiology 49, 239–243 (1978).

22. Bianchi, V. E. The anti-inflammatory effects of testosterone. J. Endocr. Soc. 3, 91–107, doi:10.1210/js.2018-00186 (2018).

23. Gubbels Bupp, M. R. Sex, the aging immune system, and chronic disease. Cell. Immunol. 294, 102–110, doi:10.1016/j.cellimm.2015.02.002 (2015).

24. Martín-Millán, M. & Castañeda, S. Estrogens, osteoarthritis and inflammation. Joint Bone Spine 80, 368–373, doi:10.1016/j.jbspin.2012.11.008 (2013).

25. Aoki, T., Yamamoto, Y., Ikenoue, T., Onishi, Y. & Fukuhara, S. Multimorbidity patterns in relation to polypharmacy and dosage frequency: a nationwide, cross-sectional study in a Japanese population. Sci. Rep. 8, 3806, doi:10.1038/s41598-018-21917-6 (2018).

26. Doos, L., Roberts, E. O., Corp, N. & Kadam, U. T. Multi-drug therapy in chronic condition multimorbidity: a systematic review. Fam. Pract. 31, 654–663, doi:10.1093/fampra/cmu056 (2014).

27. Subramanian, A. et al. A next generation connectivity map: L1000 platform and the first 1,000,000 profiles. Cell 171, 1437–1452.e1417, doi:10.1016/j.cell.2017.10.049 (2017).

28. Roman-Blas, J. A., Castañeda, S., Largo, R. & Herrero-Beaumont, G. Osteoarthritis associated with estrogen deficiency. Arthritis Res. Ther. 11, 241, doi:10.1186/ar2791 (2009).

29. de Klerk, B. M. et al. Limited evidence for a protective effect of unopposed oestrogen therapy for osteoarthritis of the hip: a systematic review. *Rheumatology (Oxford*, England) 48, 104–112, doi:10.1093/rheumatology/ken390 (2009).

30. Watt, F. E. Hand osteoarthritis, menopause and menopausal hormone therapy. Maturitas 83, 13–18, doi:10.1016/j.maturitas.2015.09.007 (2016).

31. Bar-Yehuda, S. et al. Induction of an antiinflammatory effect and prevention of cartilage damage in rat knee osteoarthritis by CF101 treatment. Arthritis Rheumatol. 60, 3061–3071, doi:10.1002/art.24817 (2009).

32. Yuan, Q., Sun, L., Li, J.-J. & An, C.-H. Elevated VEGF levels contribute to the pathogenesis of osteoarthritis. BMC Musculoskelet. Disord. 15, 437, doi:10.1186/1471-2474-15-437 (2014).

33. Nagao, M. et al. Vascular endothelial growth factor in cartilage development and osteoarthritis. Sci. Rep. 7, 13027, doi:10.1038/s41598-017-13417-w (2017).

34. Takeshita, N. et al. Alleviating effects of AS1892802, a rho kinase inhibitor, on osteoarthritic disorders in rodents. J. Pharmacol. Sci. 115, 481–489, doi:10.1254/jphs.10319FP (2011).

35. Kong, L., Wang, L., Meng, F., Cao, J. & Shen, Y. Association between smoking and risk of knee osteoarthritis: a systematic review and meta-analysis. Osteoarthr. Cartil. 25, 809–816, doi:10.1016/j.joca.2016.12.020 (2017).

36. Chou, C. H. et al. Insights into osteoarthritis progression revealed by analyses of both knee tibiofemoral compartments. Osteoarthr. Cartil. 23, 571–580, doi:10.1016/j.joca.2014.12.020 (2015).

37. Butterfield, N. C. et al. Accelerating functional gene discovery in osteoarthritis. bioRxiv, 836221, doi:10.1101/836221 (2019).

38. Bastick, A. N., Runhaar, J., Belo, J. N. & Bierma-Zeinstra, S. M. A. Prognostic factors for progression of clinical osteoarthritis of the knee: a systematic review of observational studies. Arthritis Res. Ther. 17, 152, doi:10.1186/s13075-015-0670-x (2015).

## References Online Methods

39. Steinberg, J. et al. Widespread epigenomic, transcriptomic and proteomic differences between hip osteophytic and articular chondrocytes in osteoarthritis. *Rheumatology (Oxford*, England) 57, 1481–1489, doi:10.1093/rheumatology/key101 (2018).

40. Hawtree, S., Muthana, M., Wilkinson, J. M., Akil, M. & Wilson, A. G. Histone deacetylase 1 regulates tissue destruction in rheumatoid arthritis. Hum. Mol. Genet. 24, 5367–5377, doi:10.1093/hmg/ddv258 (2015).

41. Li, H. et al. The Sequence Alignment/Map format and SAMtools. Bioinformatics 25, 2078–2079, doi:10.1093/bioinformatics/btp352 (2009).

42. Tischler, G. & Leonard, S. biobambam: tools for read pair collation based algorithms on BAM files. Source Code Biol. Med. 9, 13–13, doi:10.1186/1751-0473-9-13 (2014).

43. Andrews, S. FastQC: a quality control tool for high throughput sequence data. Available online at: http://www.bioinformatics.babraham.ac.uk/projects/fastqc. (2010).

44. Patro, R., Duggal, G., Love, M. I., Irizarry, R. A. & Kingsford, C. salmon provides fast and bias-aware quantification of transcript expression. Nat. Meth. 14, 417–419, doi:10.1038/nmeth.4197 (2017).

45. Soneson, C., Love, M. & Robinson, M. Differential analyses for RNA-seq: transcript-level estimates improve gene-level inferences [version 1; referees: 2 approved]. F1000Research 4, 1521, &lt;http://f1000r.es/6a7> (2015).

46. Purcell, S. et al. PLINK: a tool set for whole-genome association and population-based linkage analyses. Am. J. Hum. Genet. 81, 559–575, doi:10.1086/519795 (2007).

47. Chang, C. C. et al. Second-generation PLINK: rising to the challenge of larger and richer datasets. GigaScience, doi:10.1186/s13742-015-0047-8 (2015).

48. 1000 Genomes Project Consortium et al. A global reference for human genetic variation. Nature 526, 68–74, doi:10.1038/nature15393 (2015).

49. McCarthy, S. et al. A reference panel of 64,976 haplotypes for genotype imputation. Nat. Genet. 48, 1279–1283, doi:10.1038/ng.3643 (2016).

50. Das, S. et al. Next-generation genotype imputation service and methods. Nat. Genet. 48, 1284–1287, doi:10.1038/ng.3656 (2016).

51. Robinson, M. D. & Oshlack, A. A scaling normalization method for differential expression analysis of RNA-seq data. Genome Biol. 11, R25, doi:10.1186/gb-2010-11-3-r25 (2010).

52. Robinson, M. D., McCarthy, D. J. & Smyth, G. K. edgeR: a Bioconductor package for differential expression analysis of digital gene expression data. Bioinformatics 26, 139–140, doi:10.1093/bioinformatics/btp616 (2010).

53. Stegle, O., Parts, L., Piipari, M., Winn, J. & Durbin, R. Using probabilistic estimation of expression residuals (PEER) to obtain increased power and interpretability of gene expression analyses. Nat. Protoc. 7, 500–507, doi:10.1038/nprot.2011.457 (2012).

54. Ongen, H., Buil, A., Brown, A. A., Dermitzakis, E. T. & Delaneau, O. Fast and efficient QTL mapper for thousands of molecular phenotypes. Bioinformatics 32, 1479–1485, doi:10.1093/bioinformatics/btv722 (2016).

55. Sul, J. H., Han, B., Ye, C., Choi, T. & Eskin, E. Effectively identifying eQTLs from multiple tissues by combining mixed model and meta-analytic approaches. PLOS Genet. 9, e1003491, doi:10.1371/journal.pgen.1003491 (2013).

56. Han, B. & Eskin, E. Interpreting meta-analyses of genome-wide association studies. PLOS Genet. 8, e1002555, doi:10.1371/journal.pgen.1002555 (2012).

57. Giambartolomei, C. et al. Bayesian test for colocalisation between pairs of genetic association studies using summary statistics. PLOS Genet. 10, e1004383, doi:10.1371/journal.pgen.1004383 (2014).

58. Conesa, A. et al. A survey of best practices for RNA-seq data analysis. Genome Biol. 17, 13, doi:10.1186/s13059-016-0881-8 (2016).

59. Ritchie, M. E. et al. limma powers differential expression analyses for RNA-sequencing and microarray studies. Nucleic acids research 43, e47, doi:10.1093/nar/gkv007 (2015).

60. McCarthy, D. J., Chen, Y. & Smyth, G. K. Differential expression analysis of multifactor RNA-Seq experiments with respect to biological variation. Nucleic acids research 40, 4288–4297, doi:10.1093/nar/gks042 (2012).

61. Love, M. I., Huber, W. & Anders, S. Moderated estimation of fold change and dispersion for RNA-seq data with DESeq2. Genome Biol. 15, 550, doi:10.1186/s13059-014-0550-8 (2014).

62. Leek, J. T. svaseq: removing batch effects and other unwanted noise from sequencing data. Nucleic acids research 42, e161, doi:10.1093/nar/gku864 (2014).

63. Law, C. W., Chen, Y., Shi, W. & Smyth, G. K. voom: precision weights unlock linear model analysis tools for RNA-seq read counts. Genome Biol. 15, R29, doi:10.1186/gb-2014-15-2-r29 (2014).

64. Tarca, A. L. et al. A novel signaling pathway impact analysis. Bioinformatics 25, 75–82, doi:10.1093/bioinformatics/btn577 (2008).

65. Young, M. D., Wakefield, M. J., Smyth, G. K. & Oshlack, A. Gene ontology analysis for RNA-seq: accounting for selection bias. Genome Biol. 11, R14, doi:10.1186/gb-2010-11-2-r14 (2010).

66. de Leeuw, C. A., Mooij, J. M., Heskes, T. & Posthuma, D. MAGMA: Generalized gene-set analysis of GWAS data. PLOS Comput. Biol. 11, e1004219, doi:10.1371/journal.pcbi.1004219 (2015).

67. Parker, H. S. et al. Preserving biological heterogeneity with a permuted surrogate variable analysis for genomics batch correction. Bioinformatics 30, 2757–2763, doi:10.1093/bioinformatics/btu375 (2014).

68. Wilkerson, M. D. & Hayes, D. N. ConsensusClusterPlus: a class discovery tool with confidence assessments and item tracking. Bioinformatics 26, 1572–1573, doi:10.1093/bioinformatics/btq170 (2010).

69. Skarnes, W. C. et al. A conditional knockout resource for the genome-wide study of mouse gene function. Nature 474, 337–342, doi:10.1038/nature10163 (2011).

70. Glasson, S. S., Blanchet, T. J. & Morris, E. A. The surgical destabilization of the medial meniscus (DMM) model of osteoarthritis in the 129/SvEv mouse. Osteoarthr. Cartil. 15, 1061–1069, doi:10.1016/j.joca.2007.03.006 (2007).

