## Supplementary Notes for "Decoding the genomic basis of osteoarthritis"

### Supplementary Note

#### Molecular QTLs in osteoarthritis tissues

We carried out a sensitivity analysis for eQTLs and pQTLs by explicitly accounting for patient age and joint site affected (knee or hip). In each tissue, ≥89.8% of eGenes and >82% of pGenes significant in the main analysis also remained significant at 5% FDR in the sensitivity analysis, with a very strong correlation of normalized effect sizes reported for eGene and pGene variants identified across both analyses (Pearson $r \geq 0.988$, $P < 10^{-15}$).

We note that we detected a smaller number of genes with pQTLs than with eQTLs, despite a slightly higher number of tissue samples in the pQTL analysis. Generally, protein expression is influenced by RNA levels as well as translational and post-translational mechanisms, so it is possible that the regulatory effects from genetic variants is diminished. A recent study of brain samples also found fewer genes with pQTLs than with eQTLs (14.7% versus 10.7% of 5,743 genes profiled on both assays)[1]. However, comparisons between detected eQTLs and pQTLs are restricted due to differences in the technological platforms.

#### Differential protein abundance between high-grade and low-grade cartilage

As batch effects in proteomics data can be pervasive[2,3], paired samples from any patient were always assayed in the same 6-plex (cohort 1) or 10-plex (cohorts 2-4). As a sensitivity analysis, we carried out a differential abundance analysis for the proteomics data with explicit adjustment for the plexes as follows. For each protein, we calculated the log2 of normalized abundance values plus 1. We then obtained residuals from linear regression of these values on the 13 batches used in the proteomics data. These residuals were quantile normalized. We then used limma to test for differential abundance between low-grade and high-grade cartilage using the quantile normalized residuals, applying a paired design and the same method as in the main analysis. The results were highly similar to the main analysis. In particular, the correlation of protein log-fold-changes between the sensitivity and main analyses was very high (Pearson $r$=0.98 and Spearman $\rho$=0.97, both $P < 10^{-10}$). Of the 2,233 proteins with FDR<5% in the main analysis, all had a concordant direction of effect in the sensitivity analyses, and 1,796 (80.4%) were also significant at 5% FDR in the sensitivity analysis. As the low-grade and high-grade samples from the same patient were assayed in the same 10-plex, the adjustment for patient effects in the main analysis is expected to capture batch effects, as indeed confirmed by this sensitivity analysis.

#### Examples of genes with cross-omics differences between high-grade and low-grade cartilage

*COL1A2*, which showed significantly higher expression in high-grade cartilage at both omics levels, encodes the pro-alpha2 chain of type I collagen that is a prominent feature of disordered fibrocartilage repair in late osteoarthritis but is absent from intact articular cartilage[4]. *COL9A1* demonstrated lower cross-omics expression in high-grade cartilage, and encodes one of the type IX collagen alpha chains of hyaline cartilage, which is severely degenerated in osteoarthritis. *MMP19*, a matrix metalloproteinase involved in the breakdown of the extracellular matrix (ECM)[5], also demonstrated higher cross-omics expression in high-grade cartilage, in agreement with matrix disintegration processes playing a central role in cartilage degradation.

#### Pathway associations for differences between low-grade and high-grade cartilage

We identified associations with several pathways related to the extracellular matrix and signaling pathways. In particular, significant associations in the Signalling Pathway Impact Analysis (SPIA) include the activation of the "ECM-receptor interaction" in high-grade compared to low-grade cartilage (identified across RNA-level, protein-level, and cross-omics DE genes), with activations of the "Focal adhesion", "Lysosome", and "Cytokine-cytokine receptor interaction" pathways identified based on the RNA data (Figure 3c, Supplementary Table 3).

These associations agree with enrichment analyses based on Gene Ontology (GO) terms in GOseq, carried out separately for genes with higher or lower expression in high-grade cartilage compared to low-grade cartilage. Here, we identified associations with terms related to "extracellular space" for genes with higher expression on RNA-level, and separately, protein-level, and across omics levels. Genes with higher expression on RNA-level also showed significant associations with terms related to "plasma membrane", "vesicle", "cell adhesion", "cell communication", and "signaling" (Supplementary Table 3). Protein-level differentially expressed genes showed enrichments for "extracellular region" and related terms among genes with both higher and lower expression; genes with lower expression also showed associations with "small molecule metabolic process", "vesicle", "ECM structural constituent", and "cofactor binding" (Supplementary Table 3).

#### Patient stratification

A previous study using array-based gene expression data across 23 patients identified two clusters in low-grade cartilage, with differences related to inflammation[6]. Of the 9 genes validated using qRT-PCR, 6 were also present in our data, and all showed significant differences in a concordant direction between the two cartilage clusters (Supplementary Table 6). However, this study did not develop a classifier that could be applied to other data.

Another study used RNA sequencing data from low-grade cartilage of 44 knee osteoarthritis patients and applied a network-based approach to identify two clusters, with differences related to inflammation, Wnt signalling, and calcium regulation[7]. A set of 10 genes was constructed to distinguish the clusters; 7 of these passed validation in a second cohort of 16 patients. All 7 of these genes were present in our data, and 3 showed significant differences between the two cartilage clusters (all 3 in a concordant direction; Supplementary Table 6). However, the other four genes were discordant or showed a false-discovery rate of 33.5-99.8%.

We applied multi-omics factor analysis (MOFA)[8], to discover drivers of variability between samples or patients (latent factors). When analysing all tissues and omics levels together, we identified latent factors that were each predominantly related to one tissue type or omics level only. MOFA detected 6 factors that together captured 31-61% of the variance in the RNA data and up to 1% of the variance in the protein data. The first three factors captured close to 20% of the variance in at least one tissue; all three largely related to the RNA data in one of the three tissues only. The first factor is related to RNA in low-grade cartilage and strongly associated with various immune system processes (chemokines binding chemokine receptors, interleukin signalling, cytokine signalling) and the ECM (ECM organisation, collagen degradation, collagen formation, ECM proteoglycans). The second factor is related to the synovium and strongly associated with pathways related to the complement cascade, ECM organisation, ECM proteoglycans, diseases of glycosylation, collagen biosynthesis, interleukin signalling, and others. The third factor is mostly related to RNA in high-grade cartilage and has similar associations to the first factor, including all of the associations listed above for the first factor. This factor is also related to RNA in low-grade cartilage, though to a much smaller extent (capturing 19.7% variance in high-grade and 5.9% variance in low-grade cartilage), with the above terms also showing significant associations. As none of the factors showed strong relationships with the protein-level data, we repeated the analysis using RNA-level data only, with similar results.

For the comparison of consensus clustering and MOFA, we sought to capture more fine-grained within-tissue results in MOFA and thus carried out MOFA separately for each tissue. In low-grade cartilage, the first MOFA factor (i.e. the main axis of variation) explained 29% of variation in gene expression levels and demonstrated strong correspondence with the cluster assignment. In agreement with this, we found that the gene expression weights for this first factor and the log-fold-differences between clusters had very high correlation (Spearman $\rho$=0.94, $P<10^{-15}$; Supplementary Figure 5a). We would therefore expect that in each cluster, gene expression differences between samples with low versus high factor scores would mirror differences between the clusters. To test this, we compared the gene expression differences between subsets of the discrete low-grade tissue clusters at a MOFA low-grade Factor 1 threshold of 0, which corresponded most closely to the cluster assignment, with consistent assignment for 84% of patients in the first cluster (38 of 45 with score >0) and 83% of patients in the second cluster (35 out of 42 with score <0). We analysed gene expression differences between the 38 and 7 samples in cartilage-Cluster1 with MOFA Factor 1 values above and below 0, respectively. Analogously, we analysed gene expression differences between the 7 and 35 samples in cartilage-Cluster2 with MOFA low-grade Factor 1 values above and below 0, respectively. Indeed, in both comparisons, we found that 93% of genes with significant expression differences also showed significant differences between the clusters, all in the expected direction (Supplementary Figure 5b). The within-cluster gene log-fold-differences also showed high correlation with the gene expression factor weights (Spearman $\rho$>0.82, $P<10^{-15}$). These findings were also recapitulated in synovium, for which the first two identified latent factors also showed good correspondence to the division of samples into clusters (Supplementary Figure 4c). Moreover, we found high correlation between the MOFA synovium Factor 2 RNA weights and log-fold-differences between synovium clusters (Spearman correlation $\rho$=0.88, $P<10^{-15}$; Supplementary Figure 5c), as well as between MOFA synovium Factor 1 RNA weights and log-fold-differences between synovium sub-clusters (Spearman correlation $\rho$>0.94, $P<10^{-15}$; Supplementary Figure 5c).

#### Clinical profiles of molecular clusters

We confirmed that the associations between low-grade cartilage cluster assignment and sex or prescription of proton pump inhibitors were robust when sex or sex and age were explicitly accounted for (Supplementary Table 7). In fact, both associations became stronger and more significant when sex and age were accounted for (sex: OR=5.14 and *P*=0.0010 versus unadjusted OR=4.12 and *P*=0.0024; prescription of proton pump inhibitors: OR=5.16 and *P*=0.0034 versus unadjusted OR=4.21 and *P*=0.0040).

#### 7-gene classifier predicts clustering

The independent dataset of 60 samples used to validate the 7-gene classifier had been assigned into two groups by the original study, reflecting differences in complement activation and innate immunity. This group assignment corresponded to the 7-gene classifier cluster assignment for 73% of the samples (32 out of 44 samples with available data). We further considered the certainty of the original group assignment as previously published[7]. Of the 18 samples with high silhouette scores (>0.7) signifying strong cluster assignment in the original analysis, 14 had a concordant dichotomous prediction by PAMR (78%). This finding indicates strong agreement and supports the predictive potential of the 7-gene classifier in this independent dataset.

#### Mouse models of implicated genes display abnormal joint phenotypes

We generated genetically-modified mice with mutant alleles in orthologues of 7 genes with significantly different expression between low-grade and high-grade cartilage. Adult mice (post-natal day 112) from these 7 lines underwent detailed rapid-throughput joint phenotyping[9] (Methods). Bone and joint phenotypes have not been previously described for any of these lines. To quantify evidence of joint disease, we determined 18 bone and cartilage parameters in the knee joints of each mouse. We used Iodine Contrast Enhanced micro-computed tomography (ICE-μCT) to detect abnormalities in tibial articular cartilage volume and thickness, and changes in subchondral bone volume (BV; normalised to tissue volume [BV/TV]), trabecular thickness, trabecular number and mineral density. Subchondral bone mineral content was also determined by X-ray microradiography (scXRM). Damage to the articular cartilage surface was quantified using Joint Surface Replication (JSR). Development and validation of these novel methods is described in a companion paper[9]. We identified at least one abnormal joint phenotype at nominal significance ($P<0.05$) for each of the 7 genes studied (Supplementary Figures 7-8).

For three genes, there was a significantly abnormal phenotype parameter involving either the lateral (LTP) or medial tibial plateau (MTP) following Bonferroni correction for the effective number of parameters measured (Supplementary Figure 7). For the *HTRA3* (HtrA serine peptidase 3) ortholog line, we found high articular cartilage surface damage (LTP), a phenotype directly associated with osteoarthritis. Htra3 has been found to inhibit TGF-β signalling, and *Htra3* expression in mouse articular chondrocytes increases following the induction of osteoarthritis[10].

We identified low subchondral bone mineral content (LTP and MTP), low BV/TV (MTP) and low trabecular thickness (MTP) for the *MATN4* (matrilin-4) ortholog line. As validated mouse models of osteoarthritis show high subchondral bone mineral content, this suggests deletion of the *MATN4* ortholog could be protective. *Mtn4* is expressed in the developing mouse joint[11] and codes for a key component of the extracellular matrix. Mutations in the matrilin family member *MATN3* gene have been found to cause multiple epiphyseal dysplasia, a childhood skeletal dysplasia which can progress to early-onset osteoarthritis in adulthood[12].

Finally, we found low bone mineral density and increased articular cartilage thickness (MTP) for the *PDLIM1* (PDZ And LIM Domain 1) ortholog, also suggesting that loss of this gene could be protective. PDLIM1 is a cytoskeleton-associated protein that has been found to inhibit NF-κB-mediated inflammatory signaling in mice[13].

We note that there are several caveats to transferring any direction of effect between mouse models and human data. First, dysregulation of the gene in both directions could be deleterious. Moreover, the mouse experiments include a comparison of mutant and wild type mice, while the molecular comparison in patients was between high-grade and low-grade cartilage in the same individual. Moreover, the mice were assayed at 16 weeks, thus representing early-stage disease, while the human patients were in a late disease stage. However, the abnormal mouse phenotypes support the involvement of *HTRA3*, *MATN4*, and *PDLIM1* genes in osteoarthritis-relevant joint phenotypes. Further insight into the mechanisms of these genes during osteoarthritis pathogenesis can be provided in future by surgical provocation of osteoarthritis in these mice, together with inducible and/or cell-specific gene targeting to the chondrocyte or osteoblast lineages.

#### Supplementary Figures

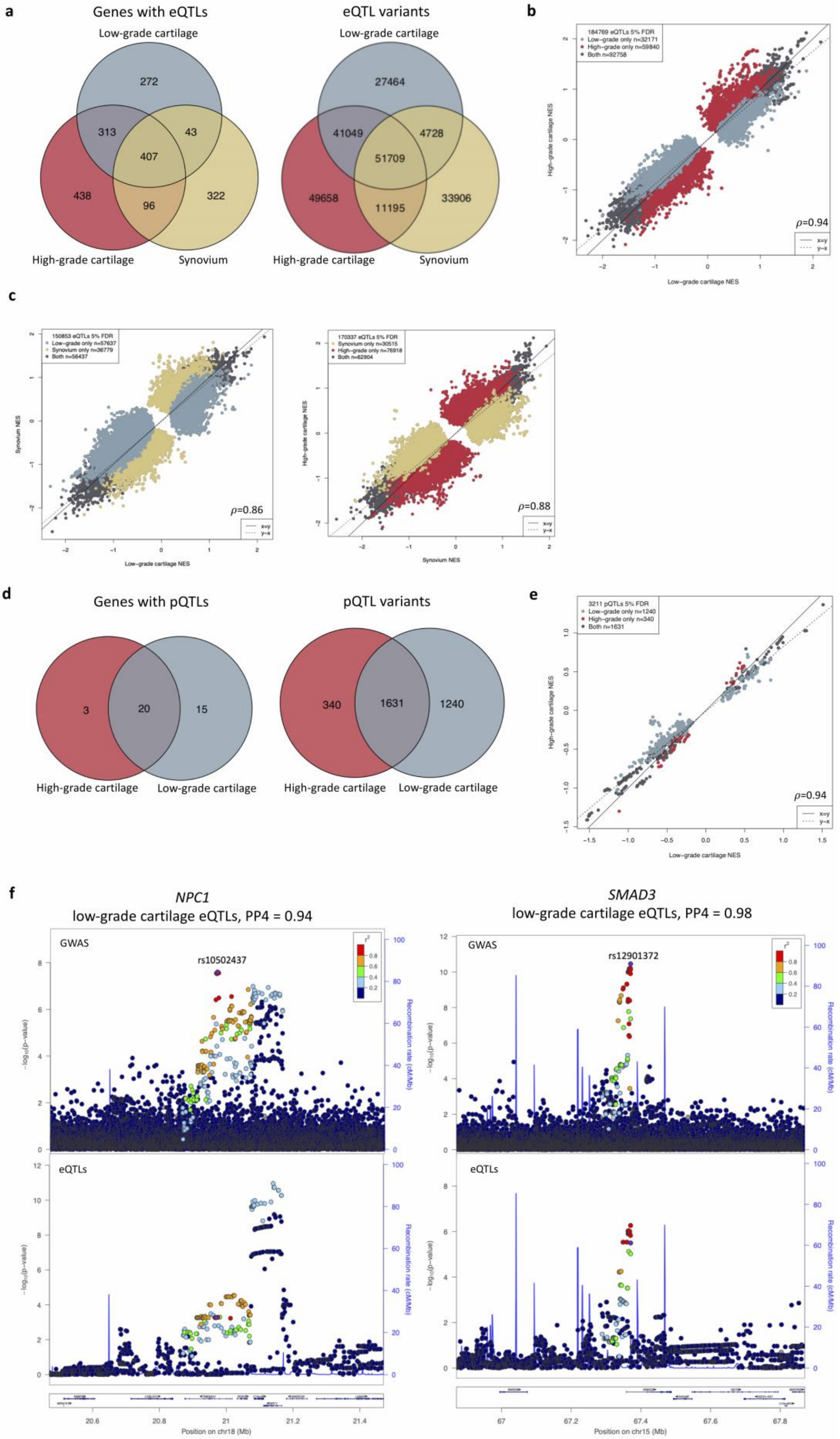
a
Genes with eQTLs
Low-grade cartilage
272
313
43
407
438
96
322
High-grade cartilage
Synovium
eQTL variants
Low-grade cartilage
27464
41049
4728
51709
49658
11195
33906
High-grade cartilage
Synovium
b
184769 eQTLs 5% FDR
Low-grade only n=32171
High-grade only n=59840
Both n=92758
High-grade cartilage NES
Low-grade cartilage NES
ρ=0.94
x=y
y=-x
c
150853 eQTLs 5% FDR
Low-grade only n=57637
Synovium only n=36779
Both n=56437
Synovium NES
Low-grade cartilage NES
ρ=0.86
170337 eQTLs 5% FDR
Synovium only n=30515
High-grade only n=76918
Both n=62904
High-grade cartilage NES
Synovium NES
ρ=0.88
d
Genes with pQTLs
3
20
15
High-grade cartilage
Low-grade cartilage
pQTL variants
340
1631
1240
High-grade cartilage
Low-grade cartilage
e
3211 pQTLs 5% FDR
Low-grade only n=1240
High-grade only n=340
Both n=1631
High-grade cartilage NES
Low-grade cartilage NES
ρ=0.94
f
NPC1
low-grade cartilage eQTLs, PP4 = 0.94
GWAS
rs10502437
eQTLs
Recombination rate (cM/Mb)
Position on chr18 (Mb)
SMAD3
low-grade cartilage eQTLs, PP4 = 0.98
GWAS
rs12901372
eQTLs
Recombination rate (cM/Mb)
Position on chr15 (Mb)

**Supplementary Figure 1. Molecular QTLs in osteoarthritis disease tissue**

a) eQTL overlap between tissues, for a total of 1,891 genes with a least one eQTL (left) and 219,709 eQTL gene-variant pairs (right). 49% of detected eQTLs are not tissue-specific.
b) High correlation of eQTL normalized effect sizes (NES) between low-grade and high-grade cartilage. Inset: Spearman correlation $\rho$=0.94 between NES effect sizes across all eQTLs.
c) High correlation of eQTL normalized effect sizes (NES) between low-grade cartilage and synovium (left), and between high-grade cartilage and synovium (right). Inset: Spearman correlation between NES effect sizes across all eQTLs.
d) pQTL overlap between tissues, for a total of 38 genes with a least one pQTL (left) and 3,211 pQTL protein-variant pairs (right).
e) High correlation of pQTL NES between low-grade and high-grade cartilage. Inset: Spearman correlation between NES effect sizes across all eQTLs.
f) Plots of GWAS and low-grade cartilage eQTL p-values for regions surrounding rs10502437 and rs12901372. For both GWAS signals, we observed colocalisation with eQTLs in both low-grade and high-grade cartilage, and the plots for high-grade cartilage are shown in Figure 2d. Each plot shows 1Mb region centered around the GWAS index SNP (purple); each point represents a genetic variant. Top panels show GWAS p-values, bottom panels QTL p-values for the indicated gene. Here and in Figure 2d, LD between variants was calculated using UK Biobank. PP4: posterior probability for colocalisation.

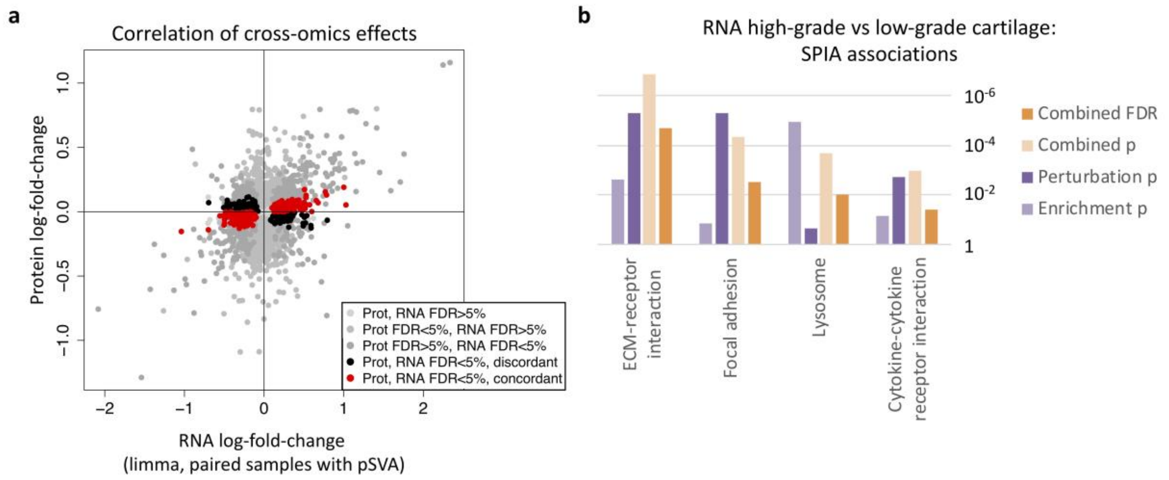

**Supplementary Figure 2. Molecular differences between low-grade and high-grade cartilage**

a) RNA- and protein-level log-fold-differences for all genes measured on both omics levels. The x-axis shows gene expression differences between low-grade and high-grade cartilage as quantified by limma in an analysis including the technical covariates as identified by pSVA as well as pairing samples from the same patients (see Methods). The genes highlighted black or red were significant on both RNA- and protein-level, see also Figure 3b. The correlation of effect sizes across all genes was also significant (Pearson *r*=0.32, *P*<10-10).

b) Signalling Pathway Impact Analysis (SPIA) identified biological pathways associated with high-grade/low-grade differences. All pathways shown are activated in high-grade compared to low-grade cartilage. Enrichment p: p-value from over-representation analysis of genes; Perturbation p: p-value for perturbation of the pathway based on gene log-fold-differences; Combined p: p-value from combining enrichment and perturbation p-values. Pathways with significant results at 5% FDR based on RNA-level changes are shown.

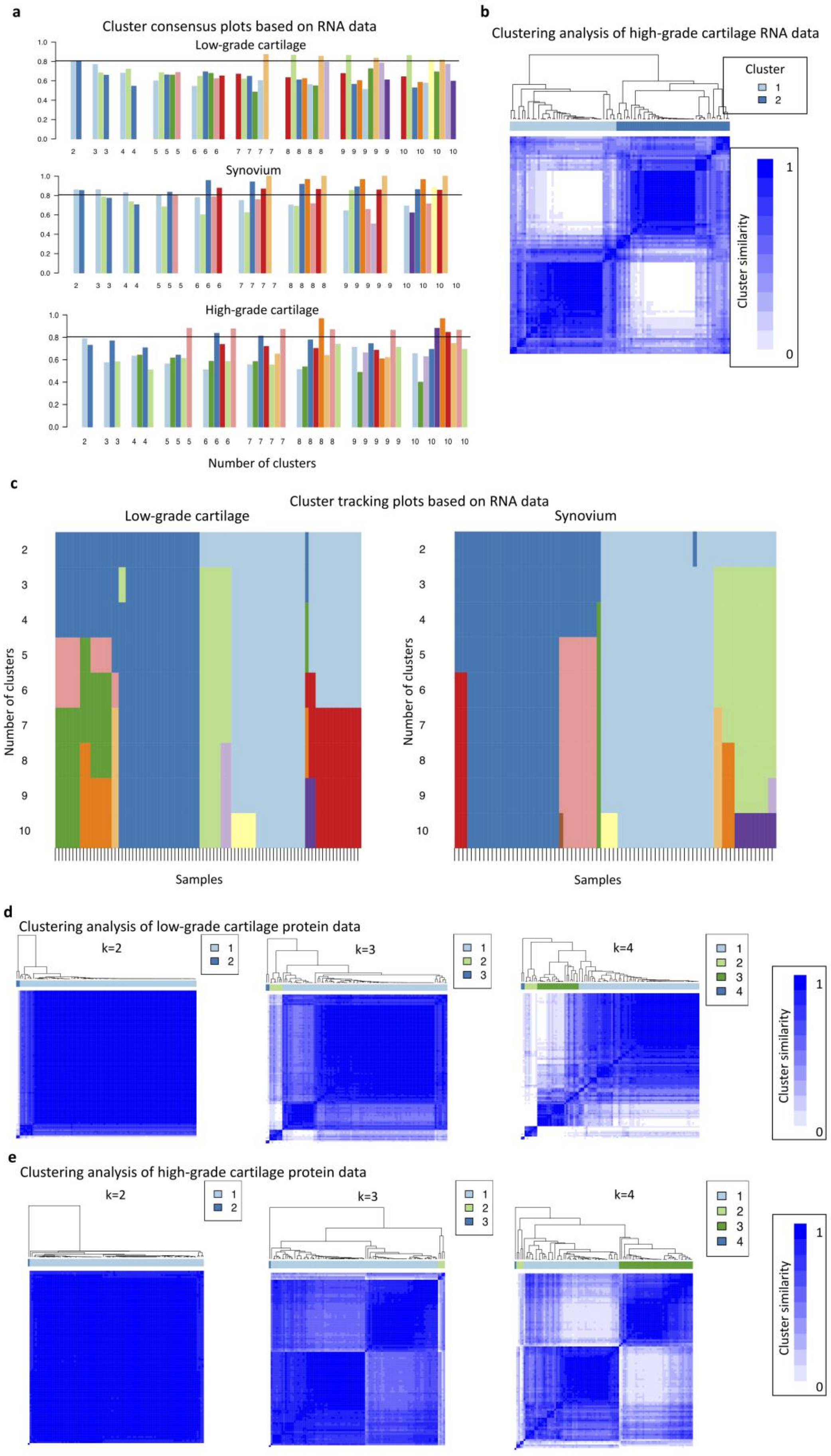

a
Cluster consensus plots based on RNA data
Low-grade cartilage
Synovium
High-grade cartilage
Number of clusters
b
Clustering analysis of high-grade cartilage RNA data
Cluster
1
2
Cluster similarity
1
0
c
Cluster tracking plots based on RNA data
Low-grade cartilage
Synovium
Number of clusters
Samples
d
Clustering analysis of low-grade cartilage protein data
k=2
k=3
k=4
Cluster similarity
e
Clustering analysis of high-grade cartilage protein data
k=2
k=3
k=4
Cluster similarity

**Supplementary Figure 3. Cluster consensus and tracking plots for the clustering analysis of samples within tissues**

a) Cluster consensus plots for clustering in low-grade cartilage, synovium, and high-grade cartilage based on RNA data. The x-axis shows the number k of clusters, the y-axis the cluster consensus value (higher values showing stronger clustering). For clustering in low-grade cartilage and synovium, but not high-grade cartilage, the cluster consensus value is above 0.8 for both clusters when k=2.
b) High-grade cartilage tissue samples from patients do not show a separation into two clusters by ConsensusCluster analysis based on RNA data (cluster consensus value <0.8 for at least one cluster).
c) Cluster tracking plots for low-grade cartilage and synovium samples based on RNA data. Each column is a sample, coloured by the cluster assignment when separating samples into k=2,…,10 clusters (k values in rows).
d) Low-grade cartilage tissue samples from patients do not show a separation into clusters by ConsensusCluster analysis based on protein data. k: number of clusters.
e) High-grade cartilage tissue samples from patients do not show a separation into clusters by ConsensusCluster analysis based on protein data. k: number of clusters.

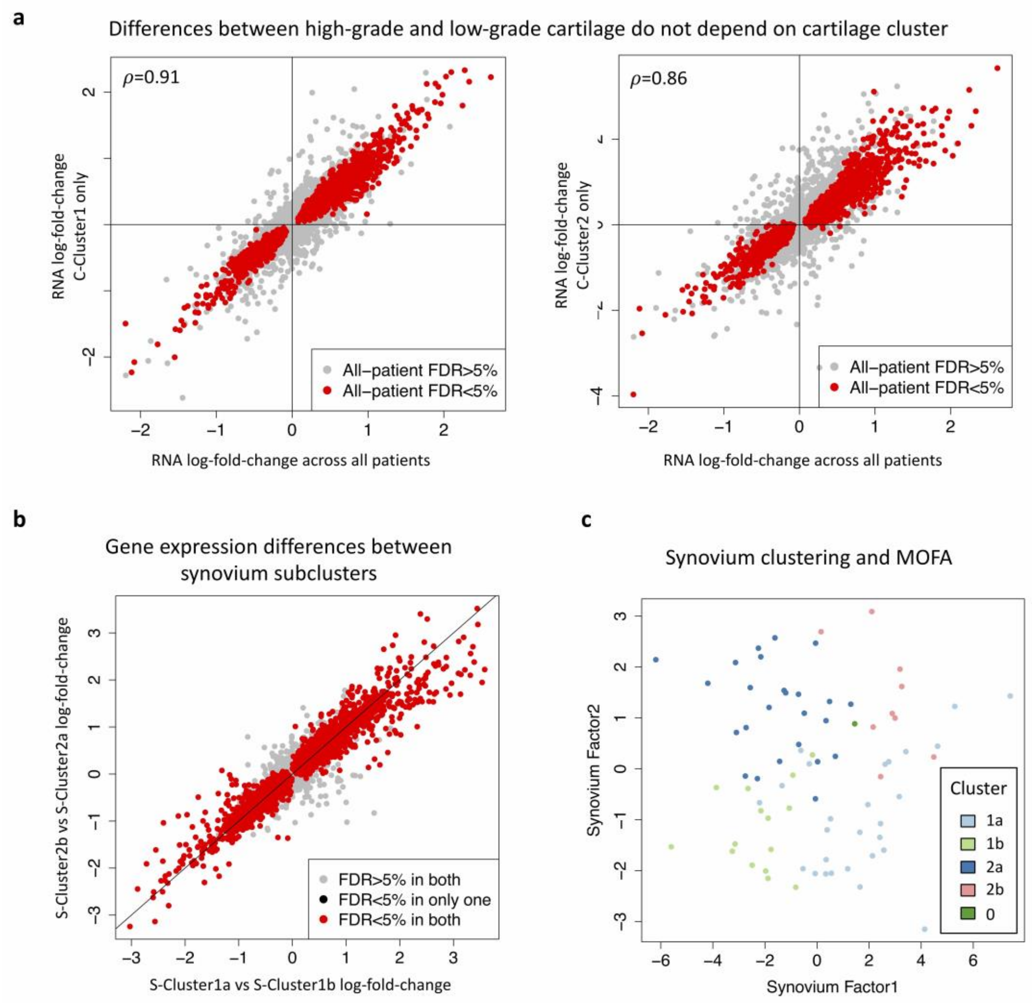

**Supplementary Figure 4. Gene expression analysis of the clusters identified in low-grade cartilage and synovium tissue and correspondence between synovium clustering and MOFA results**

a) Gene expression differences between low-grade and high-grade cartilage do not depend on the low-grade cartilage cluster. Plots show log-fold-differences for all genes based on the analysis of all patients (x-axis) versus log-fold-differences for all genes based on the analysis of all patients with low-grade cartilage in only one of the two clusters (y-axis). In each within-cluster analysis, 99% of the genes significant in the all-patient analysis had the same direction of effect. Inset: Spearman correlation of log-fold-differences, *P*<1.0x10-10.
b) Gene expression differences between the synovium sub-clusters within each cluster are highly correlated. Plot shows log-fold-differences of each gene in the comparison of sub-clusters within the larger (x-axis) and smaller (y-axis) cluster. Over 99% of the genes with significant differences between synovium-Cluster1a and synovium-Cluster1b also had directionally concordant differences between synovium-Cluster2b and synovium-Cluster2a, and over 80% were also significant at 5% FDR, and vice versa (i.e. genes with higher expression in synovium-Cluster1a compared to synovium-Cluster1b also had higher expression in synovium-Cluster2b compared to synovium-Cluster2a).

c) An analysis of synovium samples using MOFA identifies a continuous spectrum of variation between samples. This variation corresponds well to the clustering: MOFA synovium factor 1 captures differences between sub-clusters, while synovium factor 2 captures variation between clusters.

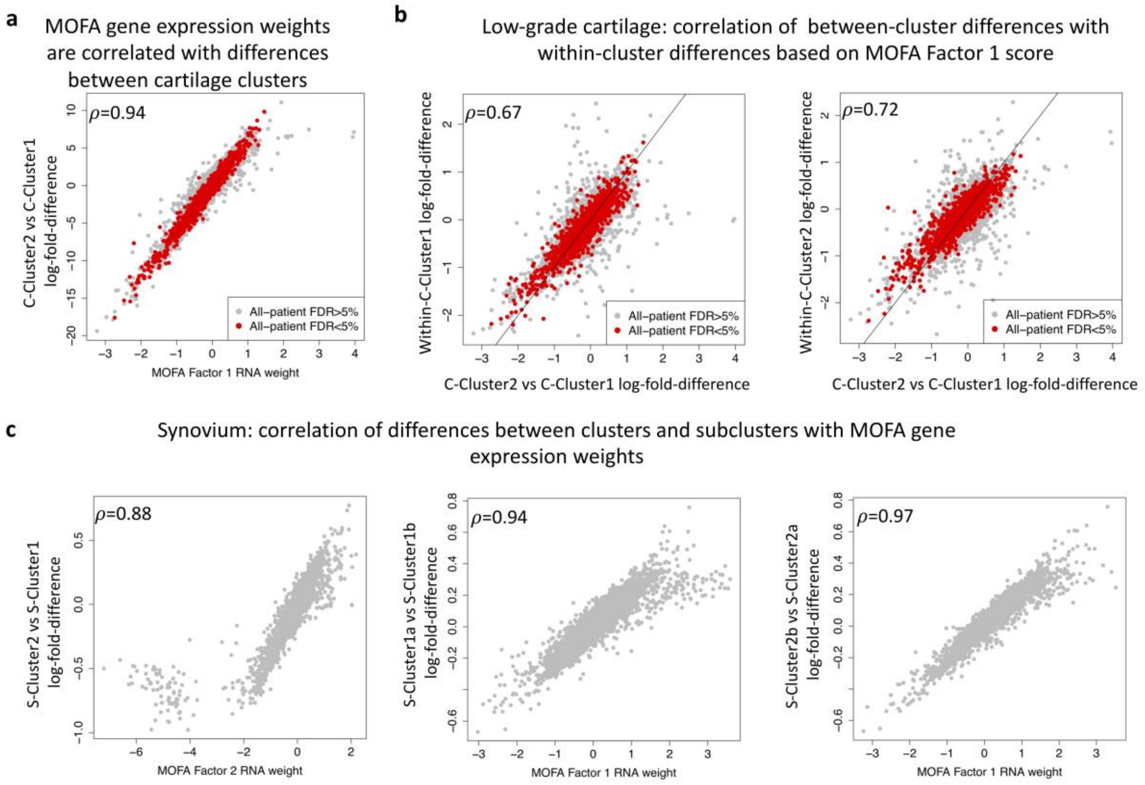

**Supplementary Figure 5. Multi-omics factor analysis (MOFA) RNA gene weights are correlated with gene expression differences between tissue clusters**

a) Correlation between MOFA low-grade cartilage Factor 1 gene weights for RNA data and gene expression differences between low-grade cartilage clusters. Inset: Spearman correlation, *P*<10-15. Genes with significant differential expression between low-grade and high-grade cartilage are coloured red.

b) Gene expression differences between low-grade cartilage samples in the same clusters, divided by MOFA low-grade cartilage factor 1 values >0 and <0, are correlated with gene expression differences between clusters. Inset: Spearman correlation, *P*<10-10. Left shows results for low-grade cartilage-Cluster1, right for cartilage-Cluster2. Genes with significant differential expression between low-grade and high-grade cartilage are coloured red.

c) Correlation between MOFA synovium Factor 1 and 2 gene weights for RNA data and gene expression differences between synovium clusters and subclusters. Inset: Spearman correlation, *P*<10-15.

C-Cluster: cartilage-Cluster.

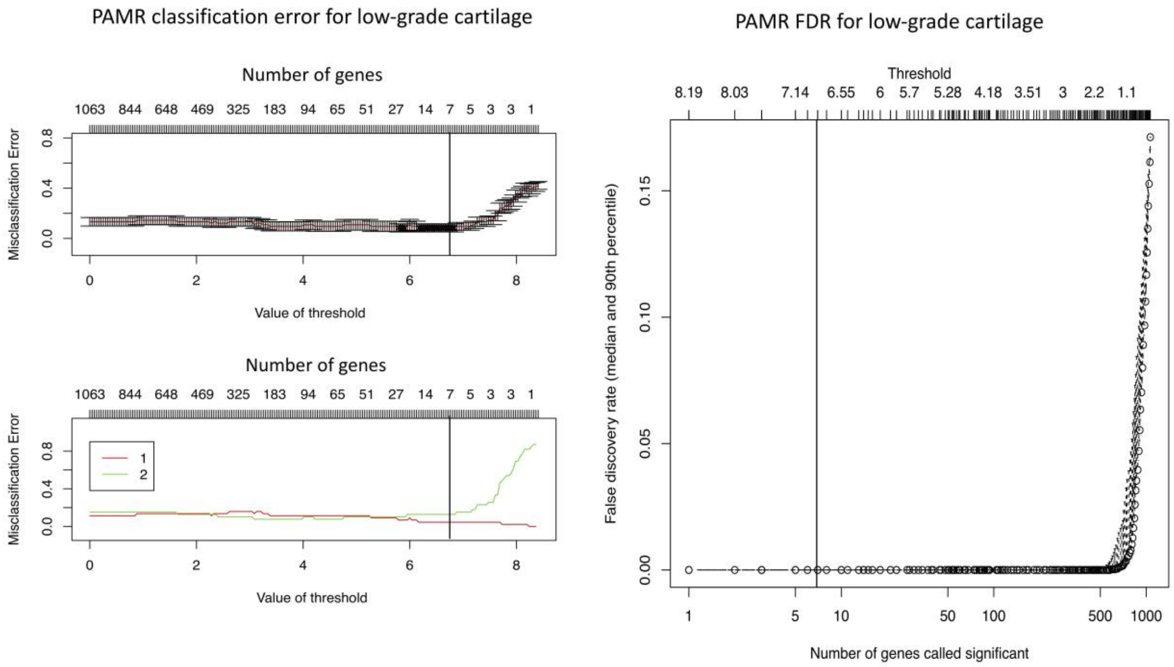

**Supplementary Figure 6. PAMR 7-gene low-grade cartilage classifier performance**
PAMR diagnostic plots for a classifier of low-grade cartilage based on RNA. Left: Sample classification error based on the PAMR internal threshold and the corresponding number of genes in the classifier. The top panel shows the overall error estimate, the bottom panel error rates separately for cartilage-Cluster1 and cartilage-Cluster2. The optimal selection as used in the paper included 7 genes and an internal threshold of 6.67 (vertical line). Right: False Discover Rate (FDR) for between-cluster differences for the genes in the classifier as calculated by PAMR. C-Cluster: low-grade cartilage-Cluster.

**a**

Abnormal mouse joint phenotypes: mutant fold-change relative to wildtype

| Gene | Articular cartilage | | | | | | | | Bone | | | | | | | | | |
|---|---|---|---|---|---|---|---|---|---|---|---|---|---|---|---|---|---|---|
| | Volume | | Median thickness | | Max thickness | | Damage area % | | Bone vol / Tissue vol | | Trabecular thickness | | Trabecular number | | BMC | | BMD | |
| | LTP | MTP | LTP | MTP | LTP | MTP | LTP | MTP | LTP | MTP | LTP | MTP | LTP | MTP | LTP | MTP | LTP | MTP |
| *Cpt1a2* $^{-/-}$ | | | | | | | | | | | | | | | 1.04 | | | |
| *Crip1* $^{-/-}$ | | | | | | | | | | | 0.85 | | 1.07 | | | | | 0.93 |
| *Clic3* $^{-/WT}$ | | | | 0.84 | 0.87 | | | | | | | | | | | | | |
| *Htra3* $^{-/-}$ | | | | | | | 3.56 | | | | | | | | | | | |
| *Sqrdl* $^{-/WT}$ | | | | | | | 3.07 | | | | | | | | | | | |
| *Matn4* $^{-/-}$ | | | | | | | | | | 0.79 | 0.81 | 0.55 | | 1.18 | 0.94 | 0.93 | | |
| *Pdlim1* $^{-/-}$ | | | | 1.21 | 1.17 | 1.16 | | | | 0.85 | 0.81 | 0.66 | 1.09 | 1.18 | | | | 0.90 |

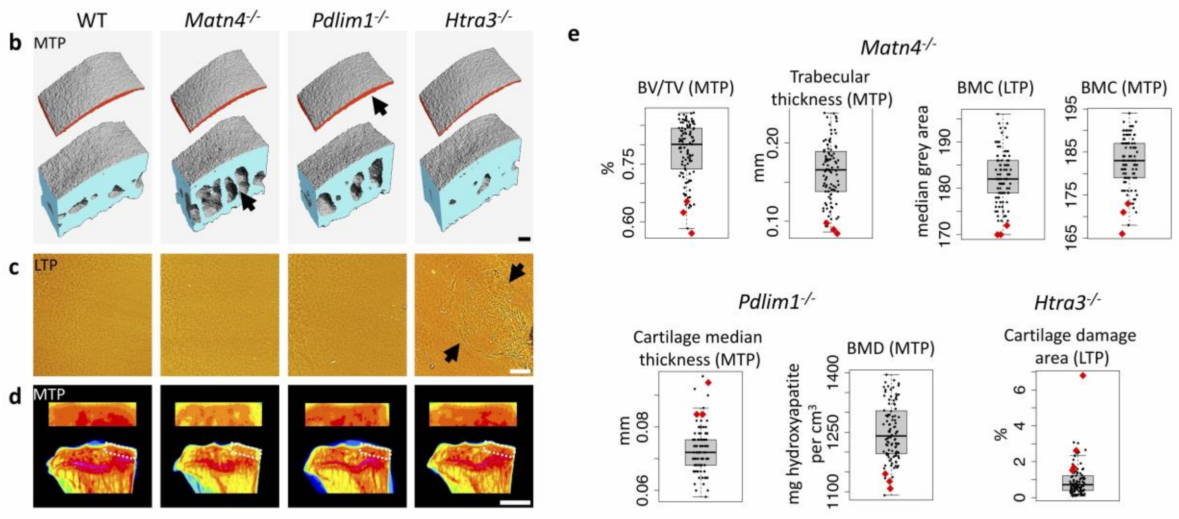

**Supplementary Figure 7. Mouse models of implicated genes display osteoarthritis-relevant abnormal joint phenotypes**

a) Overview of the abnormal mouse joint phenotypes displayed by mouse models for 7 genes with differential expression between low-grade and high-grade cartilage. For each of the genes (in rows), 9 phenotypes were assayed on the lateral tibial plateau (LTP) and medial tibial plateau (MTP). For each joint parameter (in columns), the plot shows the ratio of the mean value of each mutant strain to the mean value of the wild-type background strain, where the differences were significant (*P*<0.05). Borders around boxes show phenotype difference significant after multiple-testing correction for the effective number of tests within each line (*P*<0.00568). BMC: bone mineral content; BMD: bone mineral density; vol: volume. Technical details of each mouse mutant line and plots of all individual values for abnormal phenotypes at *P*<0.05 see Supplementary Figure 8.

b) Iodine contrast enhanced micro computed tomography (ICE-µCT) detects differences in articular cartilage volume and thickness (red volumes), and subchondral bone morphology (blue volumes); scale bar 100µm. Matn4-/- mice show decreased subchondral BV/TV (bone volume/tissue volume) and trabecular thickness (black arrow) compared to wild-type controls, whereas Pdlim1-/- mice display increased articular cartilage thickness (black arrow).

c) Joint surface replication (JSR) detects damage to the articulating surfaces of the tibial plateaux; scale bar 100um. Htra3-/- mice show increased articular cartilage surface damage (black arrows, C) compared to wild-type controls.

d) Subchondral X-ray microradiography (scXRM) detects changes in BMC within the subchondral region. White boxes represent the subchondral regions analysed; scale bar 1mm. Matn4-/- mice show decreased subchondral BMC compared to wild-type controls.

e) All abnormal phenotypes displayed by Matn4-/-, Pdlim1-/- and Htra3-/- (*P*<0.00568). The boxplots show phenotype values for 100 mice from the background strain, with error bars for the 25%-75% interquartile range. Red diamonds: phenotypes of mutant mice. BV/TV: bone volume per tissue volume.

**a**

| Gene symbol (*M.musculus*) | Targeting method | Allele name | Allele symbol |
|---|---|---|---|
| *Clic3* | Endonuclease | Clic3<em1(IMPC)Wtsi> | $Clic3^{em1/WT}$ |
| *Cpt1a* | Targeted reporter cassette | Cpt1a<tm1.2Wtsi> | $Cpt1a^{tm1.2/tm1.2}$ |
| *Crip1* | Endonuclease | Crip1<em1(IMPC)Wtsi> | $Crip1^{em1/em1}$ |
| *Htra3* | Endonuclease | Htra3<EM1> | $Htra3^{em1/em1}$ |
| *Matn4* | Endonuclease | Matn4<EM1> | $Matn4^{em1/em1}$ |
| *Pdlim1* | Endonuclease | Pdlim1<em1(IMPC)Wtsi > | $Pdlim1^{em1/em1}$ |
| *Sqrdl* | Endonuclease | Sqrdl<em1(IMPC)Wtsi> | $Sqrdl1^{em1/WT}$ |

**b**

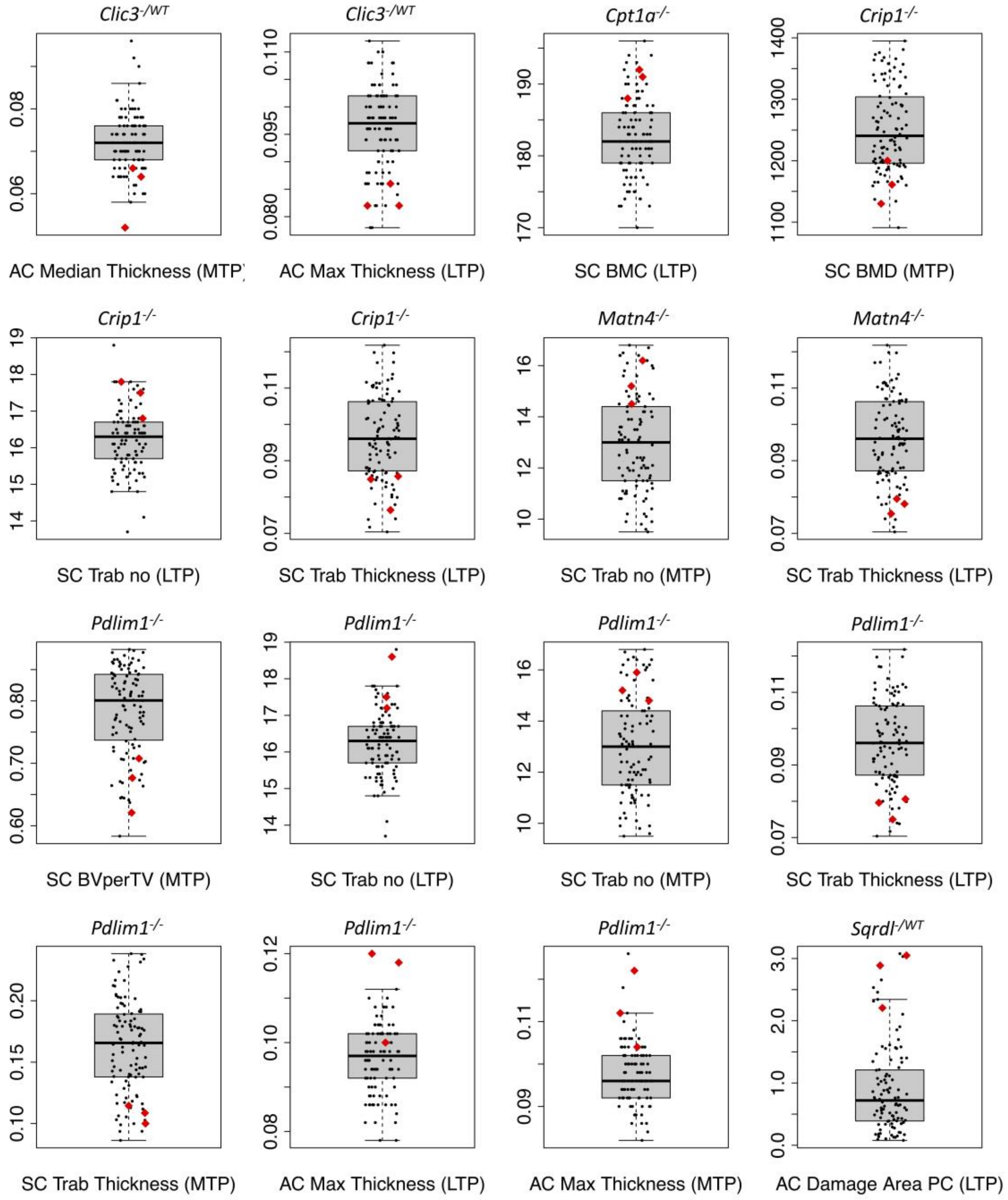

**Supplementary Figure 8. Details of mouse mutant lines and additional abnormal joint phenotypes displayed by mouse models of implicated genes**

a) Technical details for each mouse mutant line, including targeting method and allele name.

b) The figure shows all abnormal phenotypes displayed at $P<0.05$ which were not shown in the main figure. The boxplots show phenotype values for 100 mice from the background strain, with error bars for the 25%-75% interquartile range. Red diamonds: phenotypes of the mouse lines.

#### References

1 Robins, C. *et al.* Genetic control of the human brain proteome. *bioRxiv*, 816652, doi:10.1101/816652 (2019).

2 Gregori, J. *et al.* Batch effects correction improves the sensitivity of significance tests in spectral counting-based comparative discovery proteomics. *J. Proteom.* **75**, 3938-3951, doi:10.1016/j.jprot.2012.05.005 (2012).

3 Kuligowski, J. *et al.* Detection of batch effects in liquid chromatography-mass spectrometry metabolomic data using guided principal component analysis. *Talanta* **130**, 442-448, doi:10.1016/j.talanta.2014.07.031 (2014).

4 Miosge, N. *et al.* Light and electron microscopicin situhybridization of collagen type I and type II mRNA in the fibrocartilaginous tissue of late-stage osteoarthritis. *Osteoarthr. Cartil.* **6**, 278-285, doi:10.1053/joca.1998.0121 (1998).

5 Stracke, J. O. *et al.* Matrix metalloproteinases 19 and 20 cleave aggrecan and cartilage oligomeric matrix protein (COMP). *FEBS Lett.* **478**, 52-56, doi:10.1016/s0014-5793(00)01819-6 (2000).

6 Fernández-Tajes, J. *et al.* Genome-wide DNA methylation analysis of articular chondrocytes reveals a cluster of osteoarthritic patients. *Ann. Rheum. Dis.* **73**, 668-677, doi:10.1136/annrheumdis-2012-202783 (2014).

7 Soul, J. *et al.* Stratification of knee osteoarthritis: two major patient subgroups identified by genome-wide expression analysis of articular cartilage. *Ann. Rheum. Dis.* **77**, 423-423, doi:10.1136/annrheumdis-2017-212603 (2018).

8 Argelaguet, R. *et al.* Multi-Omics Factor Analysis—a framework for unsupervised integration of multi-omics data sets. *Mol. Syst. Biol.* **14**, e8124, doi:10.15252/msb.20178124 (2018).

9 Butterfield, N. C. *et al.* Accelerating functional gene discovery in osteoarthritis. *bioRxiv*, 836221, doi:10.1101/836221 (2019).

10 Tocharus, J. *et al.* Developmentally regulated expression of mouse HtrA3 and its role as an inhibitor of TGF-β signaling. *Dev. Growth Differ.* **46**, 257-274, doi:10.1111/j.1440-169X.2004.00743.x (2004).

11 Klatt, A. R., Paulsson, M. & Wagener, R. Expression of matrilins during maturation of mouse skeletal tissues. *Matrix Biol.* **21**, 289-296, doi:10.1016/S0945-053X(02)00006-9 (2002).

12 Cotterill, S. L. *et al.* Multiple epiphyseal dysplasia mutations in *MATN3* cause misfolding of the A-domain and prevent secretion of mutant matrilin-3. *Hum. Mutat.* **26**, 557-565, doi:10.1002/humu.20263 (2005).

13 Ono, R., Kaisho, T. & Tanaka, T. PDLIM1 inhibits NF-κB-mediated inflammatory signaling by sequestering the p65 subunit of NF-κB in the cytoplasm. *Sci. Rep.* **5**, 18327, doi:10.1038/srep18327 (2015).
